# The conserved ER-transmembrane protein TMEM39 coordinates with COPII to promote collagen secretion and prevent ER stress

**DOI:** 10.1101/2020.08.17.253450

**Authors:** Zhe Zhang, Shuo Luo, Guilherme Oliveira Barbosa, Meirong Bai, Thomas B. Kornberg, Dengke K. Ma

## Abstract

Dysregulation of collagen production and secretion contributes to aging and tissue fibrosis of major organs. How premature collagen proteins in the endoplasmic reticulum (ER) route as specialized cargos for secretion remains to be fully elucidated. Here, we report that TMEM39, an ER-localized transmembrane protein, regulates production and secretory cargo trafficking of procollagen. We identify the *C. elegans* ortholog TMEM-39 from an unbiased RNAi screen and show that deficiency of *tmem-39* leads to striking defects in cuticle collagen production and constitutively high ER stress response. RNAi knockdown of the *tmem-39* ortholog in *Drosophila* causes similar defects in collagen secretion from fat body cells. The cytosolic domain of human TMEM39A binds to Sec23A, a vesicle coat protein that drives collagen secretion and vesicular trafficking. TMEM-39 regulation of collagen secretion is independent of ER stress response and autophagy. We propose that roles of TMEM-39 in collagen secretion and preventing ER stress are likely evolutionarily conserved.

## Introduction

Collagen is the major molecular component of connective tissues, and the most abundant protein in animals (1). Collagen dysregulation causes many human disorders, including autoimmune diseases, brittle bone diseases (too little collagen), tissue fibrosis (too much collagen) and aging-related disorders (2–7). The multi-step biosynthesis of mature collagen by the cell is a complex process and involves procollagen gene transcription and protein translation, posttranslational modification, assembly into procollagen trimers inside the endoplasmic reticulum (ER), vesicular secretion from ER, extracellular peptide cleavage and cross-linking into collagen fibers (1, 8).

Specific mechanisms underlying the secretion of procollagen still remain poorly understood. In general, specialized intracellular vesicles defined by the coat protein complex II (COPII) transport most secreted proteins, including procollagen, from the ER to the Golgi apparatus (9, 10). Sec23, Sec24, Sec13 and Sec31 comprise COPII coat proteins, while the transport protein particle (TRAPP) complex acts a key tethering factor for COPII vesicles en route to the Golgi (11–13). Typical COPII vesicles are 60 to 80 nm in diameter, which is not sufficient for transporting procollagen trimers with up to 300 to 400 nm in length (14). In mammals, large-size COPII-coated vesicles may transport procollagen from the ER to the Golgi apparatus. TANGO1, a transmembrane protein at the ER exit site, mediates formation of specialized collagen-transporting vesicle and recruitment of procollagen (14–16). The N-terminal SH3-like domain of TANGO1 binds to the collagen chaperone HSP47 in the ER lumen, recruiting procollagens to the ER exit site (17). Its C-terminal proline-rich domain (PRD) servers as a COPII receptor by interacting with the inner shell proteins Sec23/Sec24 (18). The coil-coil domain of TANGO1 forms a stable complex with cTAE5 and SEC12, which is particularly enriched around large COPII carriers for procollagen (19). Through its membrane helices, TANGO1 organizes ER exit sites by creating a lipid diffusion barrier and an export conduit for collagen (20). While requirement of TANGO1 for secretion may depend on specific collagen types, it remains unclear whether TANGO1’s functions are broadly conserved in all animals (21, 22).

*Caenorhabditis elegans* produces over 180 collagen members that constitute the cuticle and basement membranes, encodes conserved homologs of COPII/TRAPP proteins, yet lacks apparent TANGO1 homologs (23–26). This indicates that evolutionarily conserved and TANGO1-independent mechanisms may exist in *C. elegans* to regulate procollagen secretion. From a genome-wide RNAi screen for genes affecting ER stress response, we previously identified *tmem-131* that defines a broadly conserved family of proteins important for procollagen assembly and secretion (22). Mutations in specific collagen genes, conserved COPII/TRAPP-encoding homologs, and impairment of collagen biosynthetic pathway components are known to result in a range of phenotypes including ER stress response, abnormal cuticle-associated morphology (Blister and Dumpy), and early death or growth arrest (23). *tmem-131* mutants exhibit such phenotypes typical for genes required for collagen secretion (22), while many other evolutionarily conserved genes of similar phenotype but unknown functions from our initial screen remain uncharacterized.

Here, we characterize another *C. elegans* gene *tmem-39* that encodes an ER transmembrane protein and is essential for cuticle collagen production. The deficiency of TMEM-39 protein in *C. elegans* impairs cuticle integrity and secretion of COL-19, an adult-specific cuticle collagen protein (27). We show that the *Drosophila* ortholog of *tmem-39, CG13016,* is also essential for collagen secretion. From yeast-two-hybrid (Y2H) screen, we find that the cytoplasmic loop domain of human TMEM39A binds to Sec23A, the inner-shell component of the COPII coating complex. We demonstrate that SEC-23 and other COPII proteins are also essential for collagen secretion in *C. elegans*. Our findings suggest that TMEM-39 coordinates with TMEM-131 and COPII transport machineries in the ER, and its roles in collagen secretion and preventing ER stress are likely evolutionarily conserved in multicellular animals.

## Results

### Genome-wide RNAi screen identifies *tmem-39* regulating ER stress response in *C. elegans*

We identified *D1007.5*, the sole *tmem-39* homolog in *C. elegans*, from a genome-wide RNAi screen for genes affecting the abundance of transgenic reporter *asp-17*p::GFP, which is up-regulated by temperature stress and down-regulated by ER stress (22). RNAi against *tmem-39* fully suppressed the *asp-17*p::GFP reporter expression (Fig 1A). Ensembl gene tree analysis and amino acid sequence alignment show that TMEM39 family proteins are broadly evolutionarily conserved from *C. elegans* to humans (Figs 1B and S1). A recent study reported that TMEM39A is an ER-localized transmembrane protein that regulates autophagy by controlling the trafficking of the PtdIns(4)P Phosphatase SAC1 from the ER (28). How TMEM-39 regulates ER stress response in *C. elegans* remains unknown.

**Fig 1.**
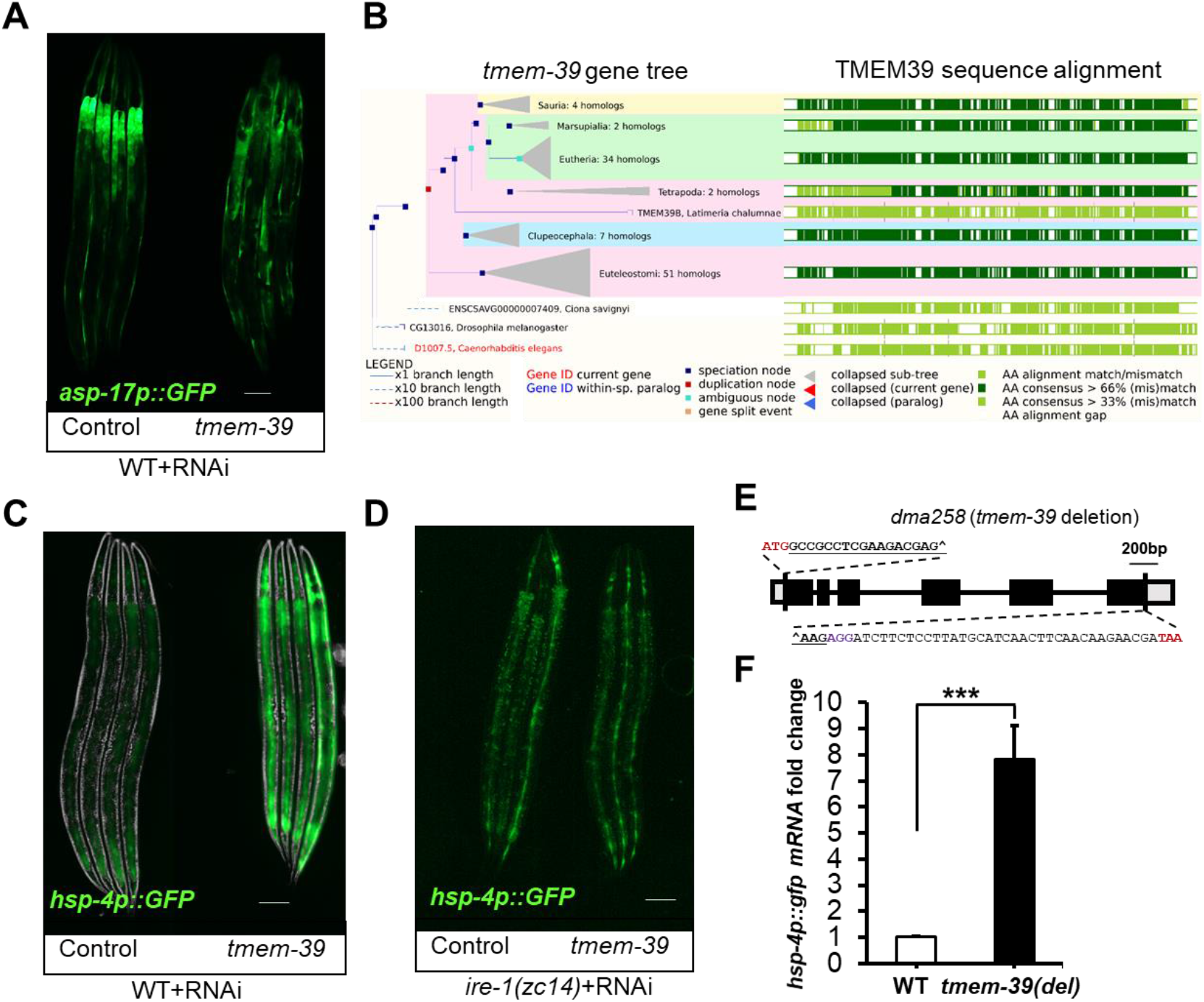
TMEM-39 regulates ER stress response in *C. elegans*. (A) Exemplar fluorescence images for *asp-17*p::GFP with control and *tmem-39* RNAi. Scale bars: 20 µm. (B) Ensembl gene tree analysis for TMEM39 protein family and amino acid sequence alignment among metazoan species (adapted from https://m.ensembl.org/). (C-D) Exemplar fluorescence and bright-field images for the UPR reporter *hsp-4*p::GFP with control and *tmem-39* RNAi in wild type (C) and *ire-1*(D) mutants. Scale bars: 20 µm. (E) Schematic of *tmem-39* gene structure with the *dma258* deletion generated by CRISPR-Cas9. Bold and underlined are sgRNA target sequence from the *tmem-3*9 regions. The location of the deleted *tmem-39* is marked with “^”. (F) qRT-PCR measurements of *hsp-4*p::GFP mRNA levels in wild-type and *dma258* mutants. ***P < 0.001 (n ≥ 3 biological replicates).

In this work, we first confirmed that RNAi against *tmem-39* in *C. elegans* caused a fully penetrant and strong up-regulation of *hsp-4*p::GFP in the hypoderm (Fig 1C). *hsp-4*p::GFP is a well-established reporter for unfolded protein response (UPR) caused by ER stress in *C. elegans* (29). Loss-of-function of IRE-1, an ER stress-sensing protein, abolished *hsp-4*p::GFP induction in *tmem-39* RNAi treated animals (Fig 1D). To verify the *tmem-39* RNAi phenotype, we used CRISPR/Cas9 to generate a *C. elegans* null allele *dma258* carrying a 2750 bp deletion of the entire coding sequence (Fig 1E and Tables S1-2). *dma258* mutants exhibited an abnormally elevated level of *hsp-4*p::GFP (Fig 1F). Besides constitutively activated *hsp-4*p::GFP transcription, TMEM-39 deficient animals by RNAi or *dma258* were small and dumpy. Together, these results suggest that TMEM-39 as an ER-localized transmembrane protein is required for maintaining normal homeostasis of ER-resident proteins.

### Loss of *tmem-39* impairs cuticle collagen secretion in *C. elegans*

To identify potential protein clients in the ER regulated by TMEM-39, we examined 24 various translational reporters of ER-resident secreted and transmembrane proteins (S2 Fig and S3 Table) and found that *tmem-39* RNAi knock-down strongly reduced abundance of the COL-19::GFP reporter (Fig 2A). COL-19 is a *C. elegans* exoskeleton collagen that is secreted by the underlying hypoderm and required for integral structure of the cuticle (23). The C-terminal GFP-tagged COL-19 reporter enables highly robust and tractable visualization of the cuticle morphology and to identify defects in the collagen production machinery (27).

**Fig 2.**
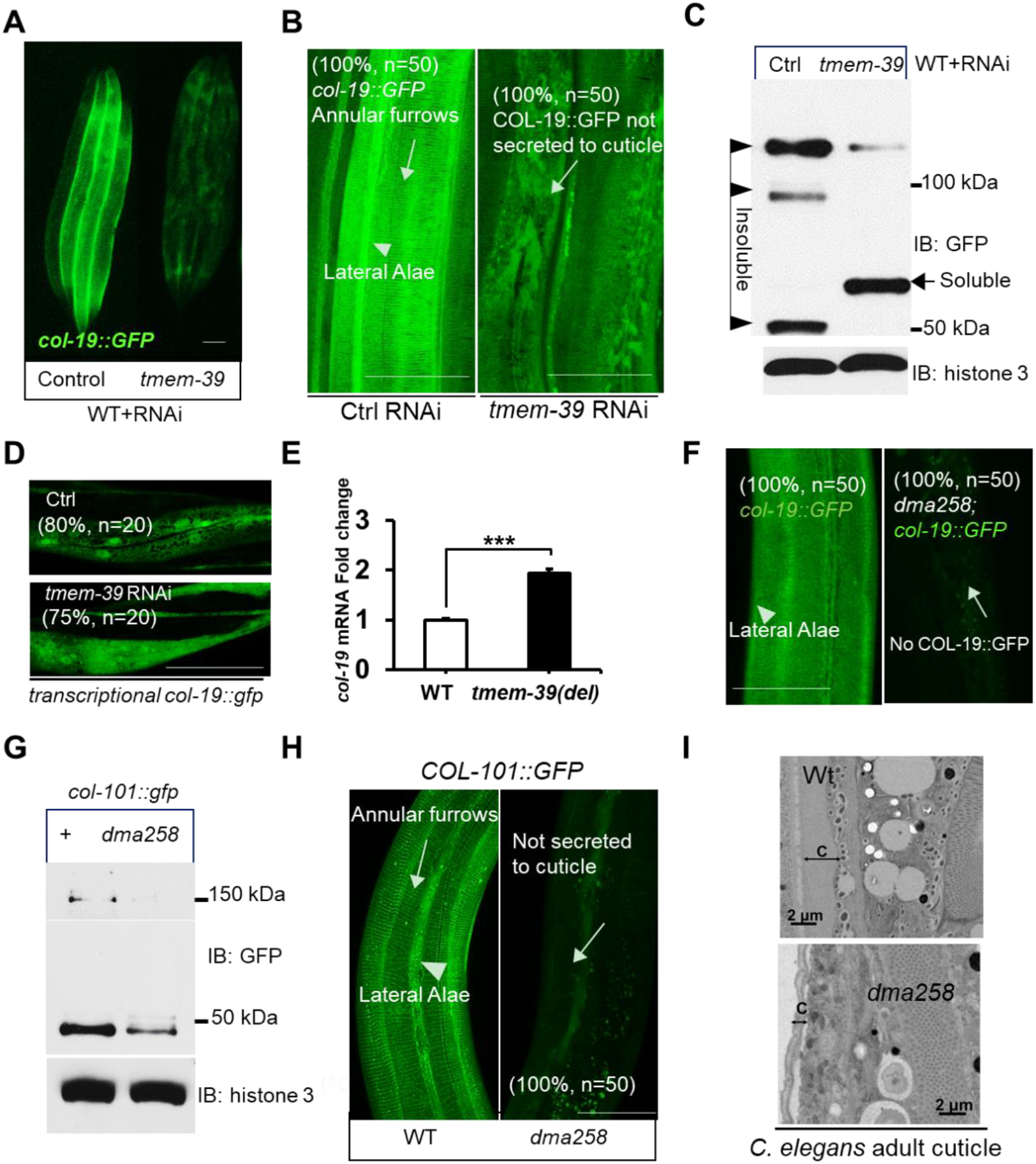
TMEM-39 is essential for collagen secretion and cuticle formation in *C. elegans*. (A) Exemplar epifluorescence image of *col-19*::GFP with control and *tmem-39* RNAi. Three to four animals were shown to indicate representative reporter expression with around 50 animals observed. (B) Exemplar confocal fluorescence images of COL-19::GFP with indicated phenotypic penetrance of control RNAi and *tmem-39* RNAi in wild-type animals. Scale bars: 20 µm. (C) Exemplar Western blot analysis of COL-19::GFP proteins from total lysates of wild type animals with control and *tmem-39* RNAi. IB, immunoblotting. The arrow indicates soluble premature monomers; triangles indicate insoluble mature monomers and cross-linked COL-19::GFP. (D) Exemplar fluorescence images of *col-19* transcriptional reporter (*col-19* promoter-driven GFP) with indicated phenotypic penetrance of control RNAi and *tmem-39* RNAi in wild-type animals, indicating no significant difference in GFP expression. Scale bars: 20 µm. (E) qRT-PCR quantification of endogenous *col-19* mRNA levels in wild-type and *dma258* mutants. ***P < 0.001 (n ≥ 3 biological replicates). (F) Exemplar confocal fluorescence images of COL-19::GFP with indicated phenotypic penetrance in wild-type and *tmem-39* mutant animals. (G-H) Exemplar images of COL-101::GFP of control RNAi and *tmem-39* RNAi in wild-type animals for Western blot analysis (G) and confocal fluorescence images (H), Scale bars: 20 µm. (I) Electron microscopy of adult *C. elegans* cross sections in wild type and *tmem-39* mutants. C, cuticle. Scale bar: 2 μm.

Using confocal microscopy to characterize the structure of hypodermal cuticle, we found that in control RNAi animals, COL-19::GFP is enriched in the hypoderm, constituting regular annular furrows and lateral alae of the cuticle (Fig 2B). In the *tmem-39* RNAi animals, COL-19::GFP appeared to be clustered in the intracellular region of hypoderm, and largely absent in the cuticle (Fig 2B). We further analyzed the abundance and composition of COL-19::GFP proteins by Western blot. Besides strong reduction of overall COL-19::GFP abundance, *tmem-39* RNAi markedly increased the soluble “premature” monomeric procollagens, while decreased the insoluble fraction of cross-linked multimers and “mature” monomers of COL-19::GFP (Fig 2C).

To examine possible involvement of *tmem-39* in collagen gene transcription, we used RNAi to knock-down *tmem-39* in animals with the *col-19*p::GFP transcriptional reporter in which *GFP* expression is driven by the promoter of *col-19*. In contrast to the striking decrease of overall COL-19::GFP protein abundance, the transcriptional activity of the *col-19* promoter was not affected by *tmem-39* (Figs 2B and 2D). We also evaluated the mRNA level of *col-19* by quantitative reverse transcription polymerase chain reaction (qRT-PCR) and found that the *dma258* mutant displayed a mild increase of *col-19* mRNA level, likely caused by compensatory feedback regulation of collagen gene transcription (Fig 2E). The *dma258* mutant fully recapitulated the *tmem-39* RNAi phenotype in defective COL-19::GFP secretion (Fig 2F).

There are two main collagen-enriched tissues in *C. elegans*, the cuticle (exoskeleton) and basement membranes (25). *tmem-39* RNAi had no effect on the production of mCherry-tagged EMB-9 (30), a Collagen IV α1 on basement membranes (S2T Fig and S3 Table). We found that loss of *tmem-39* specifically affected collagens in cuticle, as exemplified by LON-3::GFP and COL-101::GFP (Figs 2G-H and S3). Furthermore, electron microscopy (EM) analysis revealed striking reduction of cuticle thickness in *dma258* mutants than in wild type (Fig 2I). We also noticed that TMEM-39 deficient animals were small in size and dumpy, more sensitive to cuticle-disrupting osmotic stresses and developed more slowly. Taken together, these results demonstrate essential roles of TMEM-39 in collagen secretion, proper cuticle formation and preventing ER stress likely induced by premature collagen accumulation in *C. elegans*.

### Evolutionarily conserved roles of TMEM39 family proteins for collagen secretion

TMEM39 family proteins are evolutionarily conserved among multicellular animals, and the invertebrate model organisms *C. elegans* and *Drosophila* have one ortholog each, named *D1007.5* and *CG13016*, respectively (Fig 1B). We determined whether the function of TMEM39 family proteins in collagen secretion is evolutionarily conserved in *Drosophila.* We visualized collagen secretion in fat body cells of the *Lsp2> Col4a1:RFP* transgenic fly (31, 32), and generated transgenic RNAi to knock-down *Drosophila CG13016*, the sole *TMEM39* ortholog (Fig 3A). The physiological function of *Drosophila* fat body cells is to secrete collagen to the insect blood, hemolymph. Confocal microscopy analysis of COL4A1::RFP revealed that the Collagen type IV alpha 1::RFP proteins were strikingly accumulated in fat body cells of *CG13016* knock-down flies but not in control (Fig 3B). Such intracellular procollagen accumulation caused by *CG13016* RNAi indicates that the role of TMEM39 family proteins in collagen secretion is evolutionarily conserved also in *Drosophila*.

**Fig 3.**
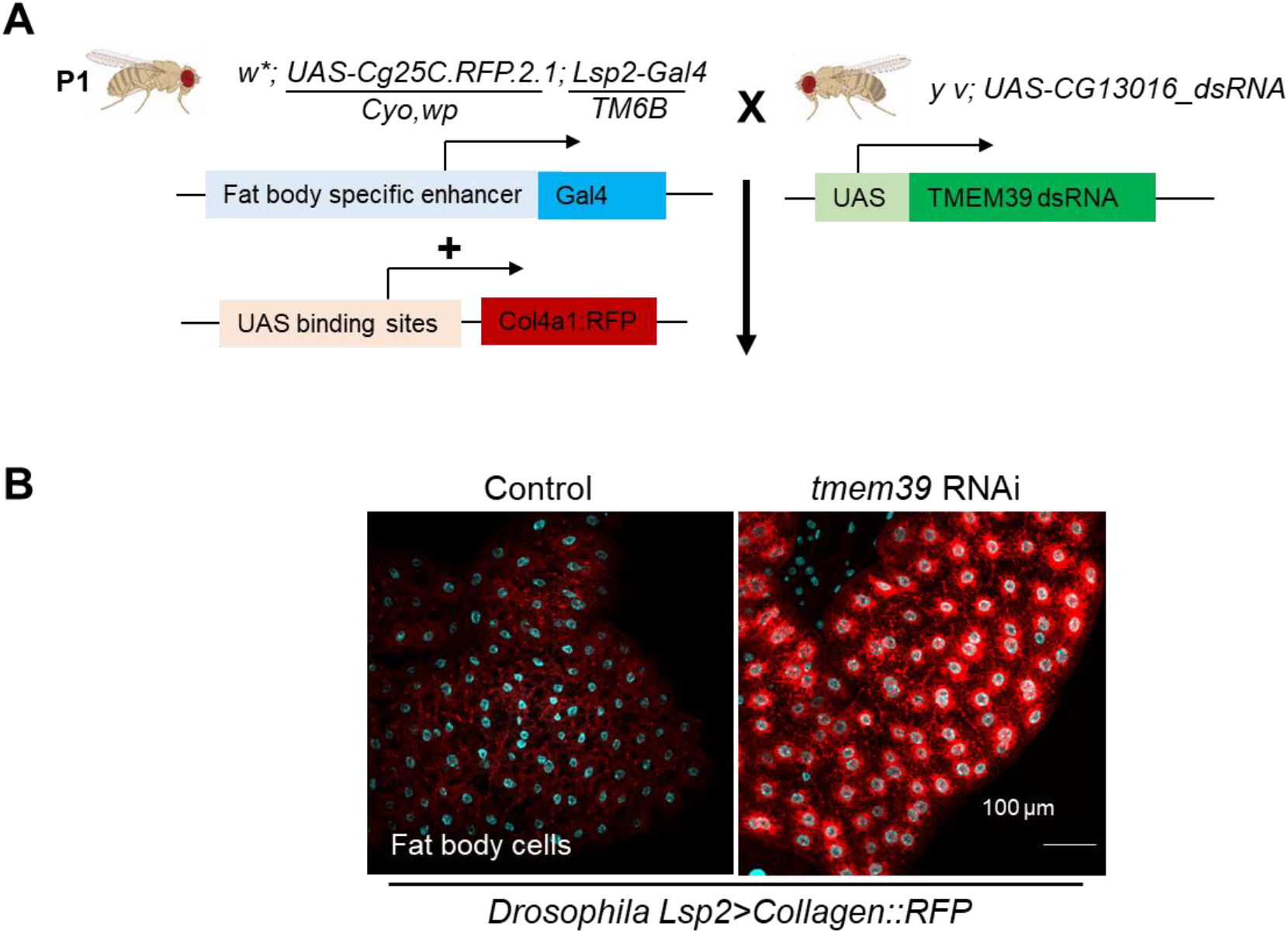
Evolutionarily conserved roles of TMEM39 family proteins for collagen secretion in *Drosophila*. (A) Schematic of generating fat body cell specific CG13016 knock-down strains in *Drosophila.* Lsp2-Gal4 specifically expresses in the fat body. Wandering third instar stage larvae were picked out for imaging analysis. The *Drosophila* images are created by BioRender.com. (B) Exemplar confocal images of transgenic *Drosophila* fat body cells showing collagen COL4A1 secretion is normal with control RNAi (left), and intracellular procollagen accumulation with *tmem39/CG13016* RNAi. scale bar, 100 µm.

Since the sequence and function of TMEM39 family proteins appear to be highly conserved, we next characterized the localization and protein interactors of human TMEM39A. The vertebrate TMEM39 family consists of two paralogs, TMEM39A and TMEM39B (33). The *TMEM39B* gene is only conserved in vertebrates, and is likely produced by the duplication of an ancestral form of *TMEM39A* (34). Consistent with a recent study (28), our confocal imaging of Hela cells transiently transfected with reporters of GFP::TMEM39A and mCherry-tagged ER markers indicates that TMEM39A localized to the ER (Fig 4A).

**Fig 4.**
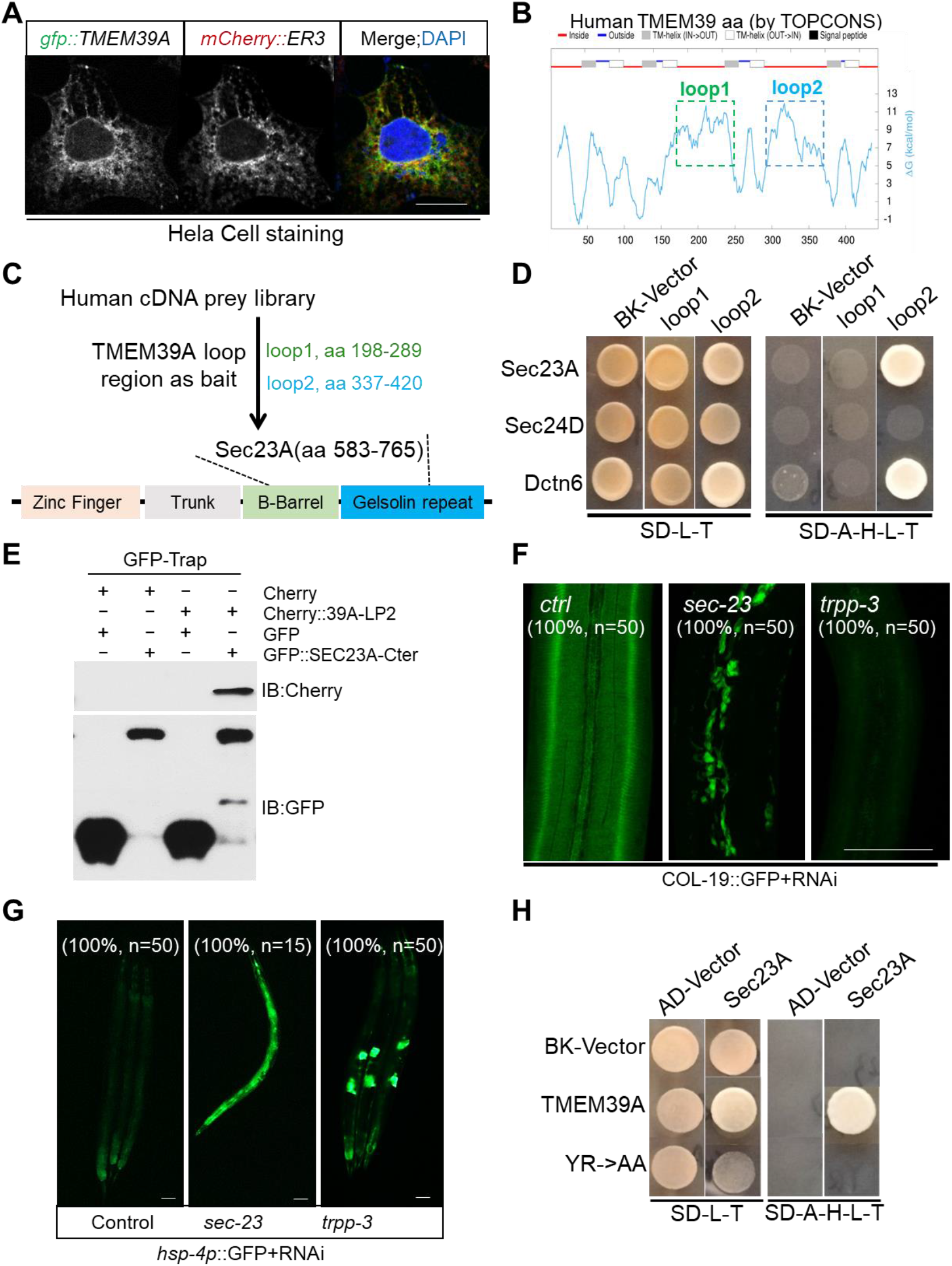
TMEM39A interacts with Sec23A to regulate collagen secretion in *C. elegans*. (A) Exemplar confocal fluorescence images of HeLa cells co-transfected with GFP-tagged TMEM39A and mCherry-tagged ER marker (ER3). Scale bars, 5 μm. (B) Schematic of human TMEM39A transmembrane domain predicted by the TOPCONS program, with cytosolic localization in red (two long cytoplasmic loop domains labeled with rectangles, loop1 in green and loop2 in blue) and ER localization in blue. (C) Schematic of Y2H screens identifying the human Sec23A C-terminal domain as a binder of the second cytoplasmic loop domain of TMEM39A. (D) Y2H assays of yeast colony growth after prey and bait vectors retransformation to verify the interaction of human Sec23A C-termini (a.a. 583-765), Sec24D full length (a.a. 1-1032) and Dctn6 full length (a.a. 1-190) with TMEM39A loop1 (a.a. 198-298) and loop2 (a.a. 337-420). (E) Coimmunoprecipitation and Western blot of mCherry-labeled TMEM39A cytoplasmic loop domain and GFP-labeled Sec23A Ct fragment in human embryonic kidney (HEK) 293 cells. Cells were transfected with expression vectors, lysed for immunoprecipitation by GFP-TRAP, and blotted by antibodies against GFP and mCherry. (F-G) Exemplar confocal fluorescence images of COL-19::GFP (F) and *hsp-4*p::GFP (G) with indicated phenotypic penetrance of wild-type with control RNAi and COPII components *sec-23* and *trpp-3* RNAi. Scale bars: 20 µm. (H) Y2H assays of yeast colony growth after prey and bait vectors retransformation to verify the interaction between human Sec23A C-termini with human wild type and YR mutant TMEM39A cytoplasmic loop domain.

### Human TMEM39A cytoplasmic loop domain interacts with Sec23A

Predicted by the TOPCONS program, TMEM-39 contains putatively eight transmembrane segments and two large cytoplasmic loops (Fig 4B). We further used the Y2H screen to search for human proteins that could interact with the conserved first loop domain (198-298 a.a.) and the second loop domain (337-420 a.a.) of TMEM39A (Fig 4C). Among the prey cDNA clones identified from the Y2H screen, Sec23A was confirmed to interact with the second loop domain of TMEM39A (Fig 4D).

The cDNA clone from the Y2H library encodes the C-terminal 583-765 a.a. of Sec23A, encompassing the Gelsolin repeat and C-terminal actin depolymerization factor-homology domain (Fig 4C). Sec23A is a core component of the COPII vesicle coating complex, which forms SEC23-SEC24 heterodimers in the inner shell of the COPII coat to select specific cargo molecules (35, 36). Mutations in human Sec23A cause an autosomal recessive disease, named Cranio-lenticulo-sutural dysplasia (CLSD) (35). The disease manifests with skeletal abnormalities, dysmorphic facial features and calvarial hypomineralization, features thought to result from defects in collagen secretion (37). Consistent with recent studies using the CoIP assay to demonstrate association between TMEM39A and Sec23A (28), we found that TMEM39A interacted with Sec23A but not Sec24D in Y2H assays (Figs 4D-E). These results indicate that the TMEM39A cytoplasmic loop domain interacts specifically with Sec23A, which forms an inner-shell heterodimer with Sec24 to drive procollagen secretion.

We next examined the loss-of-function phenotype of *sec-23*. RNAi knock-down of *sec-23*, the *C. elegans* homolog of *Sec23A*, strongly reduced COL-19::GFP secretion in the cuticle and increased its aggregation in the intracellular region of hypoderm (Fig 4F). RNAi of *sec-23* also led to strong *hsp-4*p::GFP induction, indicating constitutively activated ER stress response (Fig 4G). RNAi against genes encoding other components of COPII also recapitulated the COL-19::GFP defect and *hsp-4*p::GFP induction phenotype (Figs 4F-G, S4-5 and Table 1).

**Table 1.** RNAi of COPII-related genes for ER stress and collagen secretion phenotype analysis.

| Gene | Description | ER | collagen | Other phenotype |
| --- | --- | --- | --- | --- |
| *tmem-39* | Recruit Sec23A | +++ | ---- | Dpy, Sma, Rup |
| *tmem-131* | Recruit TRAPPC8, procollagen | +++ | ---- | Dpy, Sma, Rup |
| *sec-23* | COPII component | +++ | ---- | Lva (L1-L2) |
| *sec-24.1* | COPII component | +++ | ---- | Lva (L1-L2) |
| *sec-24.2* | COPII component | N.D. | N.D. | N.D. |
| *npp-20* | Sec13, COP II component | +++ | N.D. | Lva (L4), Rup |
| *sec-31* | COPII component | + | N.D. | N.D. |
| *sar-1* | GTPase, interact with sec-12 | +++ | -- | Lva (L4), Rup |
| *sec-12* | GTP-binding Sar1 protein | +++ | N.D. | Lva (L1-L2) |
| *rab-1* | Rab GTPase | +++ | ---- | Lva (L1-L2) |
| *trpp-3* | TRAPPIII component | ++ | ---- | N.D. |
| *trpp-6* | TRAPPIII component | N.D. | ---- | N.D. |
| *trpp-8* | TRAPPIII component | N.D. | ---- | Ste |
| *uso-1* | USO1 vesicle transport factor | +++ | N.D. | N.D. |

ER, *hsp-4p::gfp* induction for ER stress; collagen, *col-19::gfp* defect; + and -, indicate fluorescent reporter induction and reduction, respectively. N.D., no significant difference. Dpy (Dumpy), shorter and stouter than control animals at the same developmental stage; Sma (Small), shorter and thinner than control animals at the same developmental stage; Lva (larval arrest), the developmental program of the animals halts at any larval stage (L1-L4); Rup (exploded through vulva), animals are ruptured at the vulva and display an extrusion of internal organs at the site of rupture; Ste (sterile), animals are unable to produce progeny.

By amino acid sequence alignment, we identified two Tyrosine-Arginine (YR) residues in the second cytoplasmic loop domain of TMEM39A that are highly evolutionarily conserved among all examined species from invertebrates to vertebrates (S1 Fig). To test whether the conserved YR motif is important for interaction with Sec23A, we substituted the YR motif of TMEM39A into Alanine-Alanine (AA). Using Y2H assays, we found that such substitution in TMEM39A strongly attenuated its interaction with Sec23A (Fig 4H). These results show that the second cytoplasmic loop domain of TMEM39A specifically binds to the COPII inner-shell component Sec23A and its *C. elegans* homolog *sec-23* is also essential for collagen production *in vivo*.

### The collagen secretion phenotype of *tmem-39* is independent of ER stress and autophagy

We identified both *tmem-39* and *tmem-131* from the genome-wide screen for RNAi clones affecting the abundance of *asp-17*p::GFP, which is downregulated by ER stress (22). We examined collagen secretion phenotypes of other genes involved in protein modification and homeostasis in the ER identified from the *asp-17*p::GFP screen, including *ostb-1*, *nus-1*, *stt-3*, *dlst-1*, *ost-3* and *uggt-1* (Fig 5A and S4 Table). RNAi against these genes, similarly as *tmem-39* and *tmem-131*, caused marked suppression of *asp-17*p::GFP and induction of *hsp-4*p::GFP (Figs 5A-B). By contrast, RNAi knock-down of these genes did not cause COL-19::GFP collagen secretion defects (Figs 5C-D and S6A-C). We also examined additional genes that are not from the *asp-17*p::GFP screen but affect the ER stress response, including *xpb-1*, *ire-1*, *cdc-48.1*, *manf-1* and *sdf-2* in *C. elegans* (38–42). RNAi against these genes induced *hsp-4*p::GFP (Fig 5E), but did not result in collagen secretion defects (Figs 5F-G and S6D-F). These results indicate that the ER stress response is likely a consequence but not cause of cuticle secretion defects in *C. elegans* deficient in TMEM-39.

**Fig 5.**
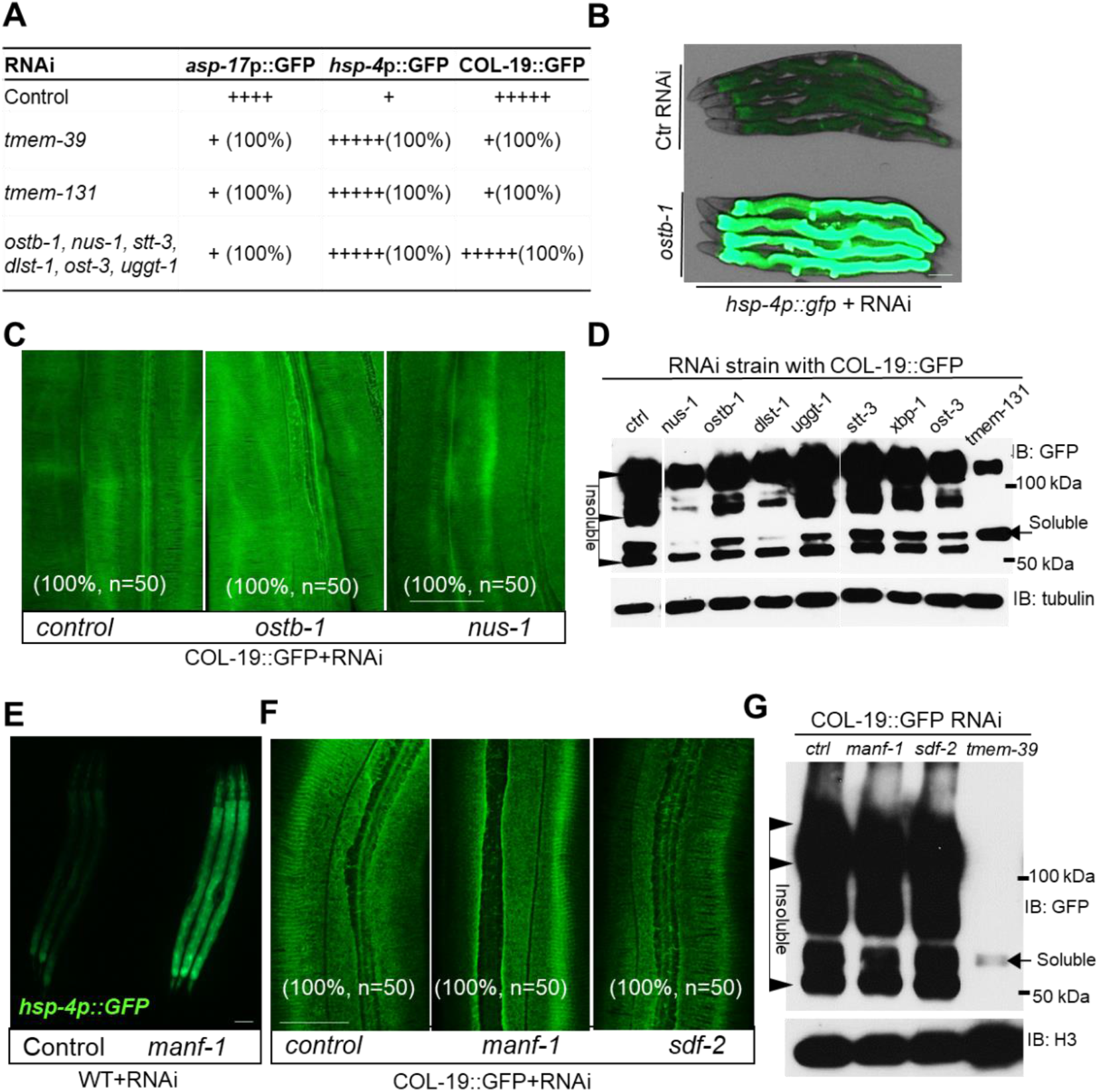
RNAi knock-down of ER stress response-related genes does not cause defects in collagen secretion. (A) Table listing ER proteostasis genes whose RNAi also suppressed *rrf-3; asp-17*p::GFP (n ≥ 20 for each group). (B) Exemplar fluorescence and bright-field images for the UPR reporter *hsp-4*p::GFP with control and *ostb-1* RNAi in wild type animals. Scale bars: 20 µm. (C-D) Exemplar confocal fluorescence images of COL-19::GFP in control RNAi and ER proteostasis gene in wild-type animals (C), scale bars: 20 µm, and Western blot analysis (D). Arrows indicate soluble premature monomers; triangles indicate insoluble mature monomers and cross-linked COL-19::GFP. (E) Exemplar fluorescence images for the UPR reporter *hsp-4*p::GFP with control and *manf-1* RNAi in wild type animals. Scale bars: 20 µm. (F-G) Exemplar confocal fluorescence images (F) and Western blot analysis (G) of COL-19::GFP with control and ER stress response gene RNAi in wild-type animals. Scale bars: 20 µm.

A recent study reported that mammalian TMEM39A regulates autophagy by controlling the trafficking of the PtdIns(4)P Phosphatase SAC1 from ER to Golgi (28). The SAC1 protein family is evolutionarily conserved among eukaryotes, while *C. elegans* has two paralogs, named SAC-1 and SAC-2 (S7 Fig). We next examined whether dysregulation of SAC-1 and autophagy might contribute to the defective collagen secretion phenotype in *tmem-39* mutants. We first confirmed that *sac-1* or *tmem-39* RNAi, but not *sac-2* RNAi, caused a marked up-regulation of the autophagy transcriptional reporter *tts-1*p::GFP (Fig 6A). *tts-1* is a long non-coding RNA that represses protein synthesis and is activated by HLH-30/TFEB, a master transcriptional regulator of autophagy (43, 44). However, *sac-1* RNAi did not affect the ER stress response reporter *hsp-4*p::GFP (Fig 6B) or COL-19::GFP (Figs 6C-D). We also examined RNAi phenotypes of *let-363,* which encodes an ortholog of human mTOR (mechanistic target of rapamycin kinase) and regulates autophagy in *C. elegans* (45, 46). Similarly as *sac-1* RNAi, *let-363* knock-down in *C. elegans* showed a marked induction of *tts-1*p::GFP but has no apparent effects on collagen secretion (Figs 6 E-G). Together, these findings indicate that roles of *C. elegans* TMEM-39 in collagen secretion are independent of ER stress response and autophagy regulation.

**Fig 6.**
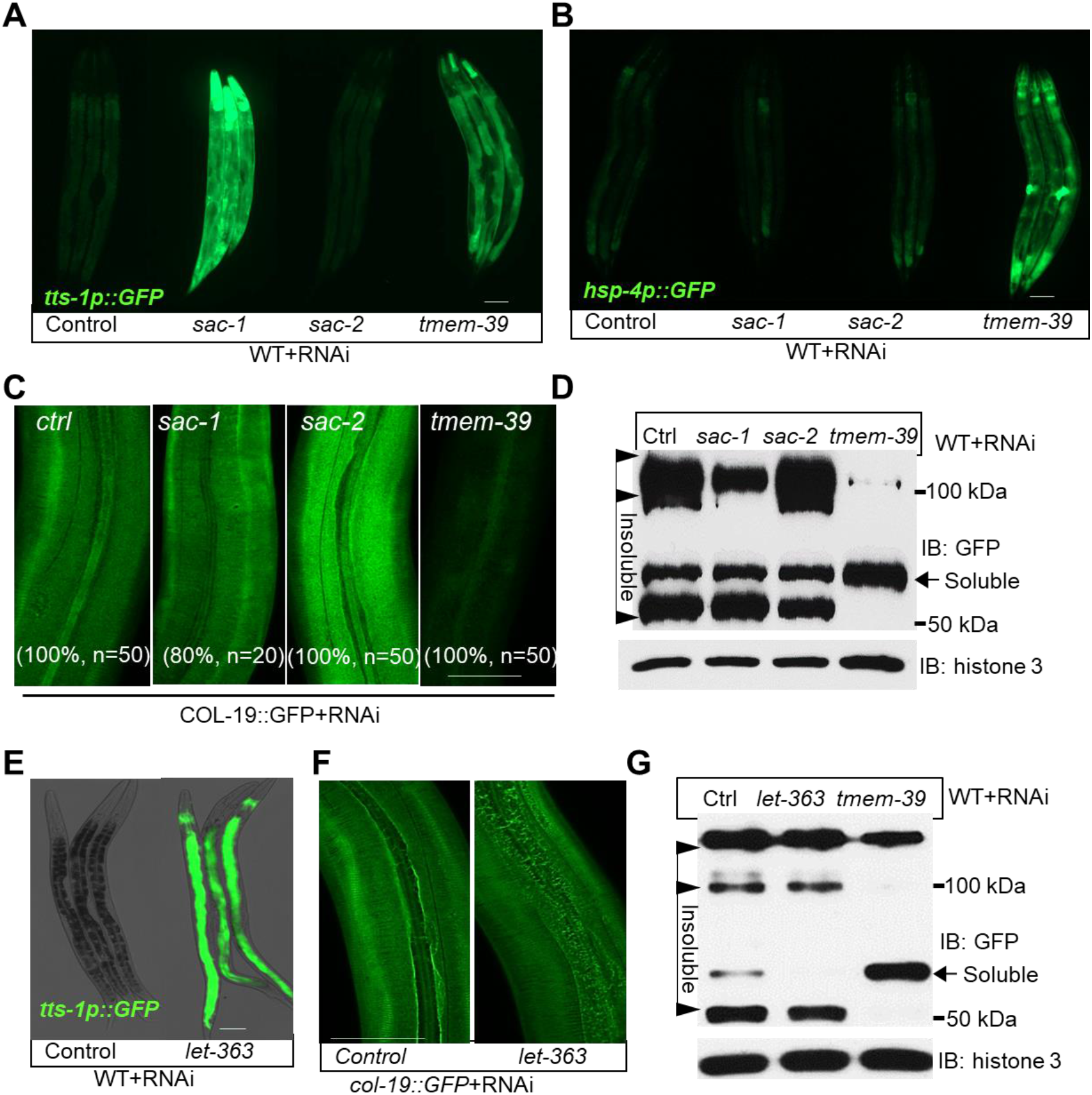
Collagen secretion is independent of ER stress and autophagy induction. (A-C) Exemplar epifluorescence images of the autophagy induction reporter *tts-1*p::GFP (A), UPR reporter *hsp-4*p::GFP (B) and COL-19::GFP (C) in *sac-1*, *sac-2* and *tmem-39* RNAi treated animals. Scale bars: 20 µm. (D) Western blot analysis of COL-19::GFP in *sac-1*, *sac-2* and *tmem-39* RNAi treated animals. Arrows indicate soluble premature monomers; triangles indicate insoluble mature monomers and cross-linked COL-19::GFP. (E) Exemplar epifluorescence images of *tts-1*p::GFP with control and *let-363* RNAi in wild type animals. Scale bars: 20 µm. (F) Exemplar confocal fluorescence images of COL-19::GFP in control and *let-363* RNAi in wild-type animals. Scale bars: 20 µm. (G) Western blot analysis of COL-19::GFP.

## Discussion

Our study identifies an ER-transmembrane protein TMEM-39 in *C. elegans* with essential roles in collagen secretion. Such roles are likely evolutionarily conserved in animals. We propose that the conserved TMEM39 cytoplasmic loop domain binds to the Sec23 component of COPII-coating complex to facilitate ER-to-Golgi procollagen transport. Phenotypic similarities of losses of TMEM-39 and TMEM-131, another ER transmembrane protein we recently identified (22), suggest that both proteins cooperate in collagen secretion by assembling premature collagen and recruiting COPII/TRAPPIII complexes for sequential ER-to-Golgi cargo transport (Fig 7).

**Fig 7.**
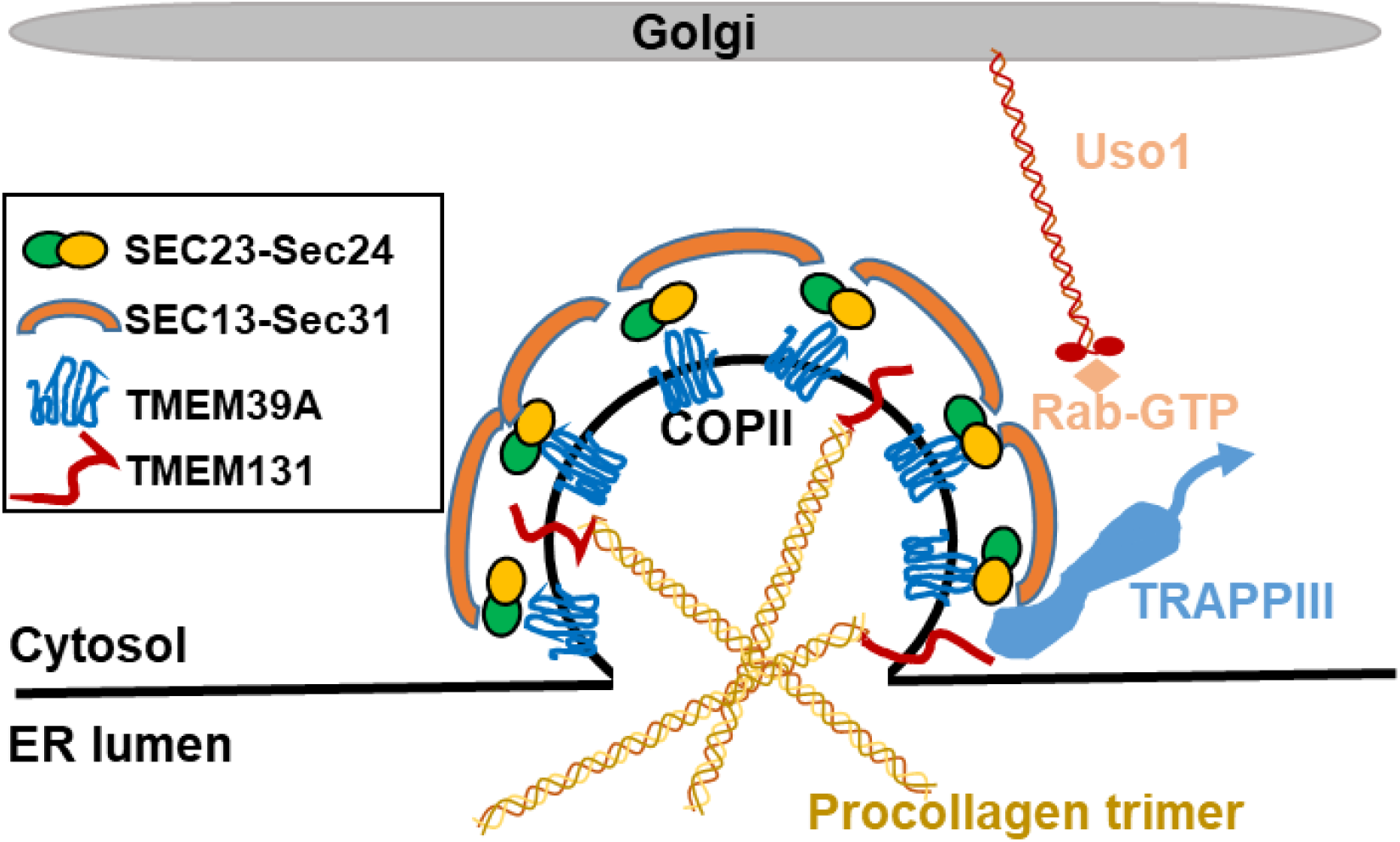
Schematic model showing TMEM39 regulation of collagen secretion. The second cytoplasmic loop domain of TMEM39A interacts with the core COPII coating component Sec23A. TMEM131 binds to COL1A2 to facilitate assembly of procollagen trimers and TRAPP III activation of Rab GTPase, in coordination with TMEM39A to promote the ER-to-Golgi transport of procollagen cargo in COPII. Uso1 interacts with the COPII vesicle to promote targeting to the Golgi apparatus.

By yeast-two-hybrid assays, we found that the TMEM39A cytoplasmic loop domain can interact with the Sec23A. RNAi knock-down of *sec-23* and most other COPII genes recapitulated the *tmem-39* loss-of-function phenotypes in constitutively high ER stress response, defective collagen secretion and sensitivity to osmolality stress in *C. elegans* (Table 1). We also noticed that RNAi knock-down of many COPII related genes, such as *sec-23, sec-24.1, npp-20, sar-1, sec-12, rab-5 and trpp-8* caused more severe phenotypes than *tmem-39* RNAi, leading to lethality or developmental arrest that prevent collagen phenotype analysis (Table 1). However, treatment with these RNAi starting from L4-stage for animals transferred from normal conditions to RNAi led to robust COL-19::GFP phenotype (Figs 4F and S4). Shorter duration of RNAi treatment may explain milder collagen defective phenotype for *sec-31*, *npp-20* and *sec-12* (Figs S4E, H and I). Compared with most COPII-related genes, *tmem-39* null mutants exhibit similar collagen secretion defects but are nonetheless viable, supporting the notion that TMEM-39 acts with COPII in collagen secretion but may have more specialized roles in facilitating secretion of specific client proteins including collagen COL-19 and the PtdIns(4)P Phosphatase SAC1 (28).

Recent work showed that TMEM39A facilitates the ER-to-Golgi transport of SAC1 and regulates autophagosome formation (28). We found that RNAi knock-down of autophagy related genes, such as *sac-1* and *let-363*, caused autophagy induction but did not affect the ER stress response or collagen secretion (Fig 6). Genes identified from the *asp-17*p::GFP screen that regulate the ER stress response also did not affect collagen secretion (S4 Table), further supporting the notion that roles of TMEM-39 in collagen secretion are independent of ER stress response and autophagy.

Besides Sec23A, additional interactors were identified from Y2H screens with the TMEM39A cytoplasmic domain as bait. We verified the interaction between full-length DCTN6 (1-190 a.a.) and TMEM39A (337-420 a.a.) (Fig 4D). DCTN6 is a subunit of the dynactin protein complex (47) that acts as an essential cofactor of the cytoplasmic dynein motor to transport a variety of cargos and organelles along the microtubule-based cytoskeleton (48, 49). In mammalian cells, ER-to-Golgi transport proceeds by cargo assembly into COPII-coated ER export sites (ERES) followed by vesicular/tubular transport along microtubule tracks toward the Golgi in a dynein/dynactin-dependent manner (50). Sec23p directly interacts with the dynactin complex (50), indicating that TMEM39A may participate in a Sec23/DCTN6 complex to facilitate COPII coat assembly and subsequent dynein/dynactin-dependent transport. Test of this hypothetic model and determination of the underlying mechanism in relation to TMEM131’s role in collagen secretion await further investigations.

Mammalian genomes encode two TMEM39 family proteins, TMEM39A and TMEM39B. *TMEM39A* is a susceptibility locus associated with various autoimmune diseases and highly up-regulated in brain tumors (33, 51). TMEM39B was recently found to interact with the SARS-CoV-2 ORF9C protein, which localizes to ER-derived vesicles (52, 53). It remains unknown whether *TMEM39A* and *TMEM39B* exhibit functional redundancy in physiological collagen secretion or pathological processes in human diseases. With single *tmem39* orthologue for each, model organisms *C. elegans* and *Drosophila* may continue to provide insights into functions and mechanisms of action of this protein family. Future elucidation of evolutionarily conserved roles of mammalian TMEM39 proteins in physiological and pathological processes may lead to therapeutic targets and strategies for treating diseases associated with this protein family in humans.

## Materials and Methods

### Worm Strains

The Bristol N2 strain was used as the wild type strain, and genotypes of other strains used are as follow: *zcIs4* [*hsp-4*p::GFP] V, *ire-1(zc14)* II; *zcIs4* V, *dmaIs10* [*asp-17*p::GFP; *unc-54*p::mCherry] X, *nIs617* [*tts-1p*::GFP, *unc-54*p::mCherry] and *kaIs12* [*col-19*::GFP], *tmem-39(dma258)* and *tmem-39(dma312)* I. Transgenic strains *dmaEx169* [*rpl-28*p::T19D2.1::mCherry; *unc-122*p::GFP], *dmaEx153* [*rpl-28*p::Y73E7A.8::mCherry; *unc-122*p::GFP], and *dmaEx152* [*rpl-28*p::F23H12.5::mCherry; *unc-122*p::GFP] were generated as extrachromosomal arrays as described (54).

The precise *tmem-39(dma258)* knock-out strain was generated by CRISPR/Cas9 methods (55, 56). Primer sequences are listed in Supporting Information Tables S1-2. Translational fluorescent reporters used by *tmem-39* RNAi knock-down to identify a phenotype include: *bcIs39* [*lim-7*p::*ced-1*::GFP+*lin-15*(+)], *caIs618* [*eff-1*p::*eff-1*::GFP], *dnSi4* [*gna-1*p::GFP + Cbr-*unc-119*(+)], *juEx1111* [*spon-1*::vGFP], *lrp-1(ku156)eqIs1* [*lrp-1*p::*lrp-1*::GFP] I; *rrf-3(pk1426)* II, *muIs49* [*egl-20*::GFP+*unc-22*(+)], *nIs590* [*fat-7*p::*fat-7*::GFP], *nuIs26* [*cat-1*::GFP], *osIs60* [*unc-5*4p::*mig-23*::GFP; *unc-119*(+)], *osIs66* [*myo-3*p::eGFP::*wrk-1*], *sqIs11* [*lgg-1*p::mCherry::GFP::*lgg-1*+*rol-6*(+)], *osIs77* [*unc-54*p::RFP::SP12;*unc-119*(+)], *pwIs503* [*vha-6*p::*mans*::GFP+Cbr-*unc-119*(+)], *qyIs44* [*emb-9*p::*emb-9*::mCherry], *rhIs23* [GFP::*him-4*], *veIs13 [col-19::GFP + rol-6(+)] V; let-7(mn112) unc-3(e151) X; mgEx725 [lin-4::let-7 + ttx-3::RFP], vkEx1243* [*nhx-2*p::ubiquitin-V::mCherry+*myo-2*p::GFP], *vkEx1256* [*nhx-2*p::*cpl-1*::YFP], *vkEx1260* [*nhx-2*p::*cpl-1*::YFP], *vkEx1879* [*nhx-2*p::*cpl-1*(W32A Y35A)::YFP] and *xnIs96* [*hmr-1*p::*hmr-1*::GFP].

### Worm maintenance

*C. elegans* strains were maintained in standard nematode growth medium (NGM) plates with seeded *E. coli* at 20 °C (57). Worm stages were synchronized by bleaching the gravid adults, and bacterial feeding-induced RNAi knock-down was performed as previously described (58). For RNAi colonies that show lethality or larvae arrest phenotypes, around 20-30 P0 L4 animals were transferred from normal NGM plates to RNAi plates, and grew for 2-3 days to observe the P0 phenotype.

### Imaging

Digital automated epifluorescence microscopes (EVOS, Life Technologies) and SPE confocal microscope (Leica) were used to obtain fluorescence images. Animals at the same stage were randomly picked from the plate, and transferred to a 4% agar pad with 10 mM sodium azide and 1 mM levamisole in M9 solution (31742-250MG, Sigma-Aldrich) on a slide for imaging. Identical setting and conditions were used to compare experimental groups with controls.

### Co-immunoprecipitation

HEK293T cells were transfected with the indicated plasmids, following the instruction of TurboFect Transfection Reagent (Thermo Fisher Scientific, R0531). After transfection for 48 hr, cells were lysed on ice for 30 min in cell lysis buffer (Cell signaling, 9803) with protease inhibitor cocktail (SIGMA 11836153001). After centrifugation at 13,000 rpm for 15 mins at 4 °C, supernatants were collected and precleaned by control magnetic beads (bmab-20, ChromoTek) for 30 mins at 4 °C, and followed by immunoprecipitation with GFP-Trap agarose beads (gtma-10, ChromoTek) for 2 hr at 4 °C. After washing with 1XPBS for 4 times and cell lysis buffer for 1 time at 4 degree, the bound proteins were eluted with 1xSDS Laemmli Sample Buffer with 10% β-mercaptoethanol and analyzed by immunoblotting.

### Western blot analysis of proteins

Animals at the same stage from the control and experiment groups were picked (N>30) into 20 µL Laemmli Sample Buffer with 10% β-mercaptoethanol and lysed directly for Western blot analysis. Protein samples were run with 15% SDS-PAGE (Bio-Rad, 4561084), and then transferred to the nitrocellulose membrane (Bio-Rad, 1620167). The membranes were blotted by antibodies against GFP (A02020, Abbkine), mCherry (Invitrogen, M11217), Tubulin (Sigma, T5168) and H3 (Abcam, ab1791).

### Quantitative RT-PCR

Worm total RNA was extracted by following the protocol of Quick-RNA MiniPrep kit (Zymo Research, R1055). cDNA was reverse transcribed by the reverse transcriptase mix kit (BioTools, B24408). Using SYBR Green Supermix (Thermo Fisher Scientific, FERK1081), the real-time qPCR was performed on the Roche LightCycler96 (Roche, 05815916001) system. Ct values of specific genes were normalized to the *C. elegans* housekeeping gene *act-1* levels. Results were presented as fold changes to respective references. Statistical significance was determined with t-test, using GraphPad Prism 7. Primer sequences are listed in Supporting information S2 Table.

### *Drosophila* experiments

Fly strains included: UAS-Cg25C:RFP.2.1/CyO; Lsp2-Gal4/TM6B, and UAS-CG13016_dsRNA (Vienna Drosophila Resource Center ID# 42509/GD). Lsp2-Gal4 is specifically expressed in the fat body cells. Flies expressing Collagen:RFP in fat body were crossed to either wild type or UAS-CG13016_dsRNA flies. Wandering-stage third instar larvae were picked out. Fat body was dissected and fixed in 4% PFA, stained with DAPI, and mounted for imaging by confocal microscopy.

### Yeast-two-hybrid assay

The cDNA coding sequences of the first and second cytoplasmic loop domain of human TMEM39A were cloned into the pGBKT7 vector and screened against a normalized universal human cDNA library (Clontech, 630481), following instruction of the Matchmaker® Gold Yeast Two-Hybrid System (Clontech, 630489). Verification of positive colonies was achieved by co-transforming wild-type or YR-mutant TMEM39A loop domain (in pGBKT7 Vector) with genes of interest (in pGADT7 Vector) following the instruction of YeastMaker™ Yeast Transformation System 2 (Clontech, 630439) as well as plasmids from re-cloned cDNA.

### Fluorescent imaging of Hela cells

Hela cells were seeded in 24-well plates with cover glass, each with three replicates (Fisher Scientific, 22293232). Cells were transiently transfected with GFP-tagged human *TMEM39A* full-length cDNA in the FUGW plasmid backbone, and the ER localization marker mCherry-ER-3 (Addgene: 55041) for 2 days. After 1xPBS washing for once, cells were treated by 4% formaldehyde solution for 10 mins. With 1xPBS washing for three times, cells were treated with 0.2% Triton X-100 in 1xPBS solution for 15 mins. Following 1xPBS washing for three times, the cover slide with cell samples was sealed on the microscope slide with Fluoroshield Mounting Medium with DAPI (Thermo Fisher Scientific, NC0200574) for imaging by confocal microscopy.

## Acknowledgements

We thank the *Caenorhabditis* Genetics Center for *C. elegans* strains and acknowledge funding support from NIH grant R01GM117461, Pew Scholar Award, Shurl & Kay Curci Foundation Faculty Scholars Program of the Innovative Genomics Institute and Packard Fellowship in Science and Engineering (to D.K.M). We thank Drs. Hong Zhang, Kamran Atabai and Matthew Shoulders for discussion and Dr. Jose Pastor-Pareja for the transgenic *Drosophila* line: w*; UAS-Cg25C.RFP.2.1. The authors declare that they have no competing interests and all participated experimental design, execution or data interpretation and analysis. Z.Z. and D.K.M wrote the paper.

## Supporting Information

**S1 to S4 Figures.**

**S1 to S4 Tables.**

**S1 Fig.**
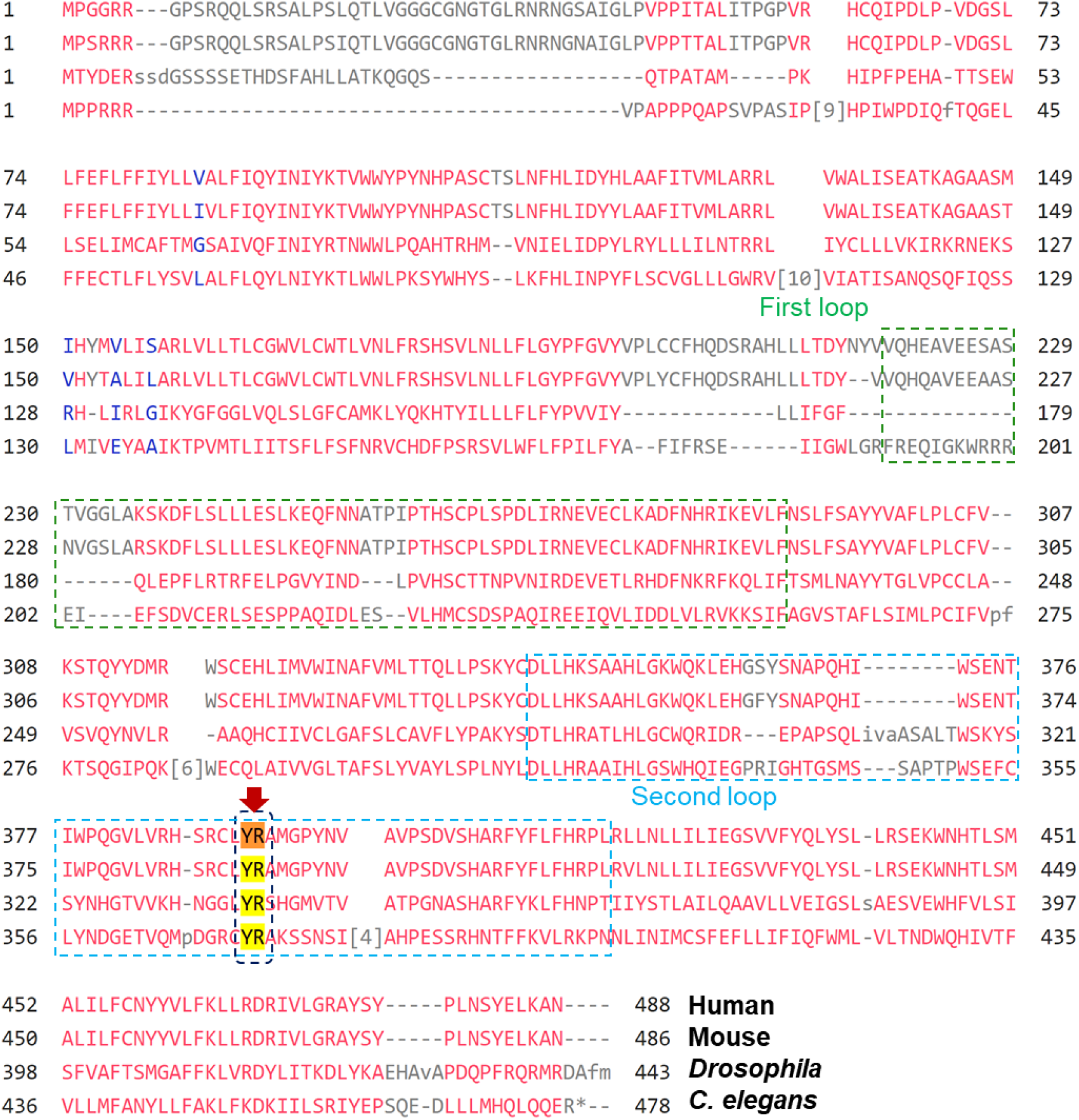
Evolutionarily conservation of TMEM39A protein sequences among different species. Multiple sequence alignment of TMEM39A from major representative animal species (by COBALT program), with conserved YR residues indicated in the second cytoplasmic loop domains.

**S2 Fig.**
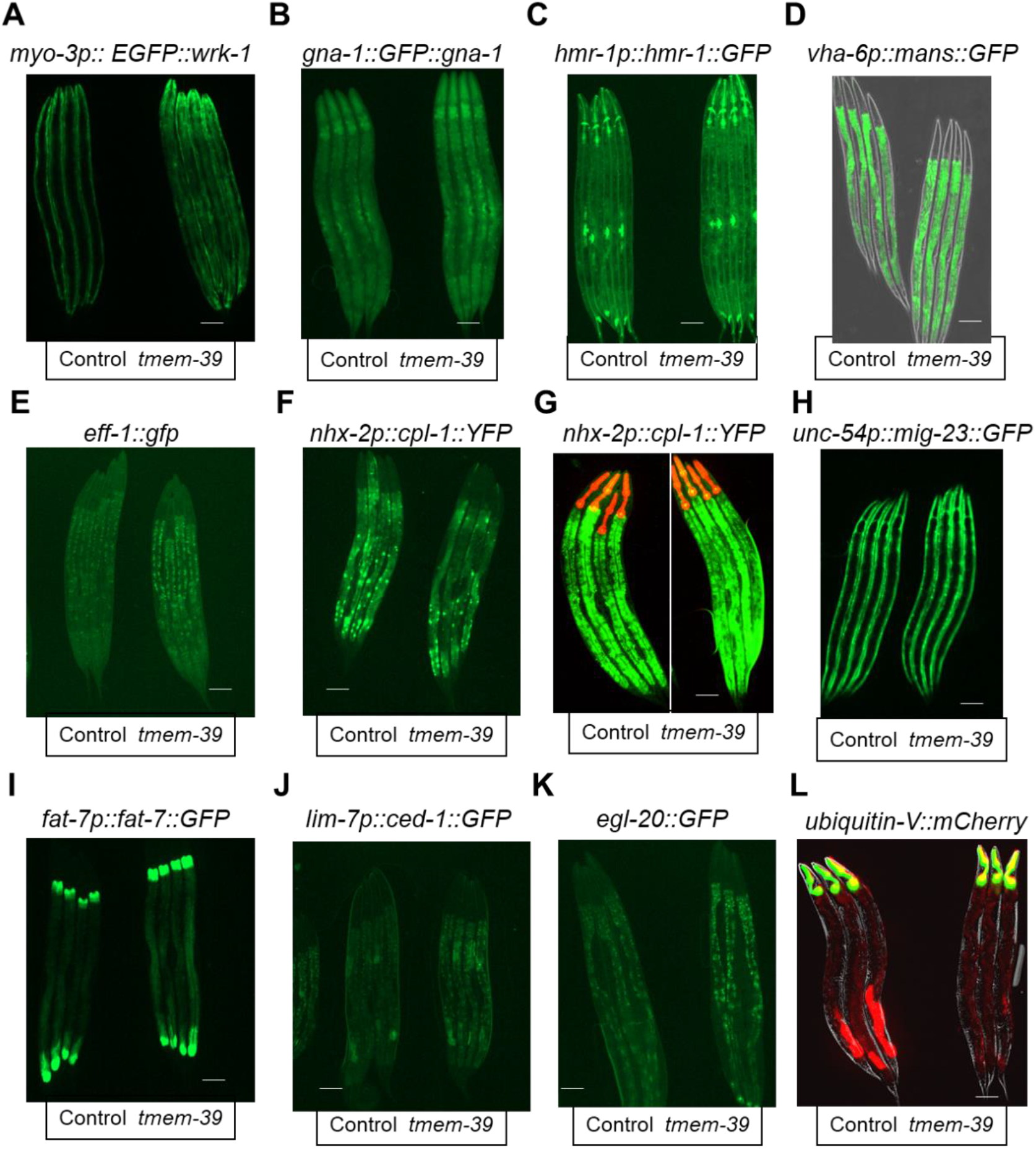
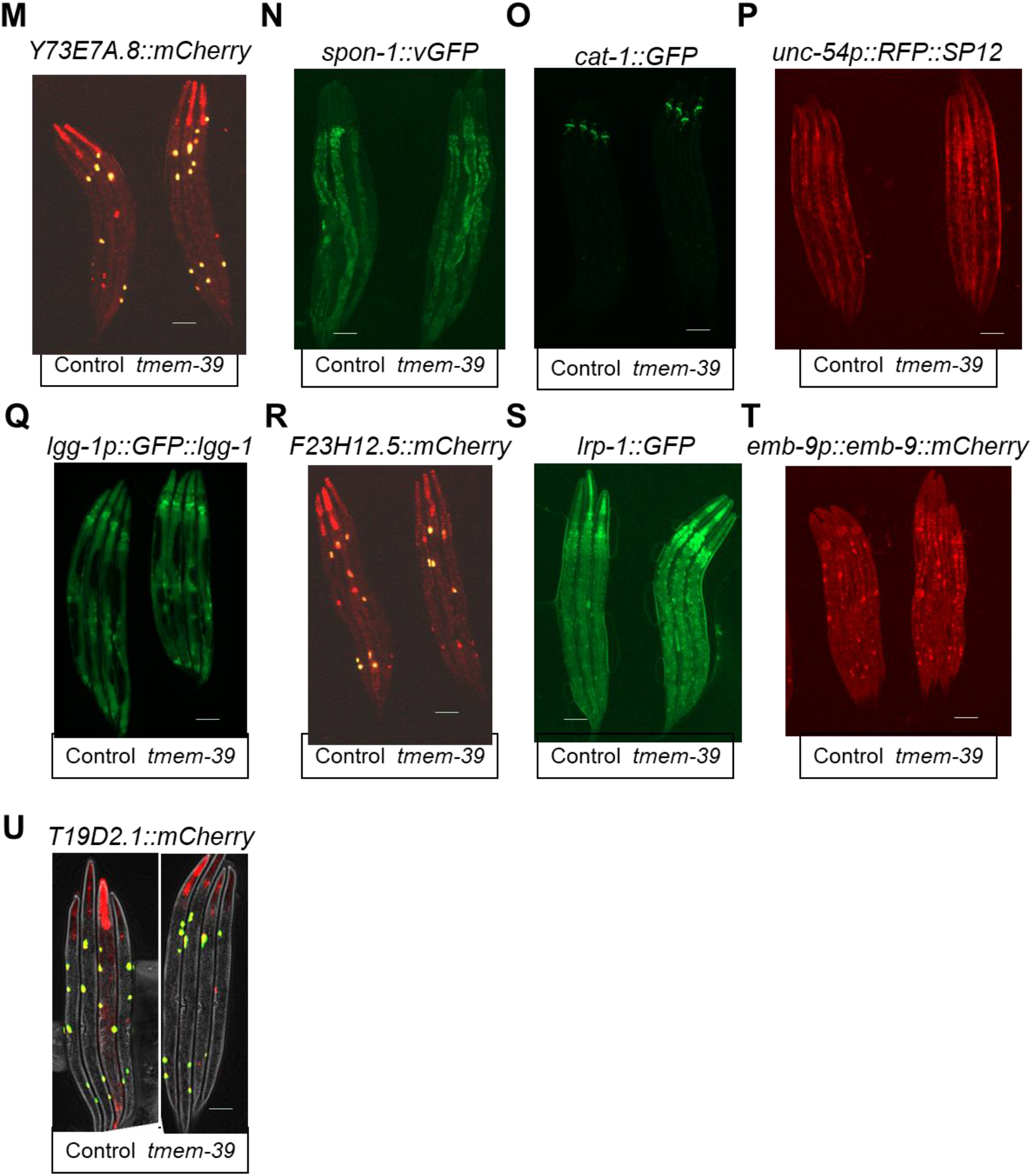
*tmem-39* RNAi knock-down for screen of phenotypic defects of different translational fluorescent reporters. (A-V) Exemplar fluorescence images showing translational reporters for (A) *wrk-1*, (B*) gna-1*, (C) *hmr-1*, (D) *mans*, (E) *eff-1*, (F-G) *cpl-1*, (H) *mig-23*, (I) *fat-7*, (J) *ced-1*, (K) *egl-20*, (L) *ubiquitin-V*, (M) *Y73E7A.8*, (N) *spon-1*, (O) *cat-1*, (P) *SP12*, (Q) *lgg-1*, (R) *F23H12.5*, (S) *lrp-1*, (T) *emb-9* and (U) *T19D2.1* in wild-type animals by control and *tmem-39* RNAi at 20 °C (n = 3-4 for each reporters). Scale bars: 20 µm.

**S3 Fig.**
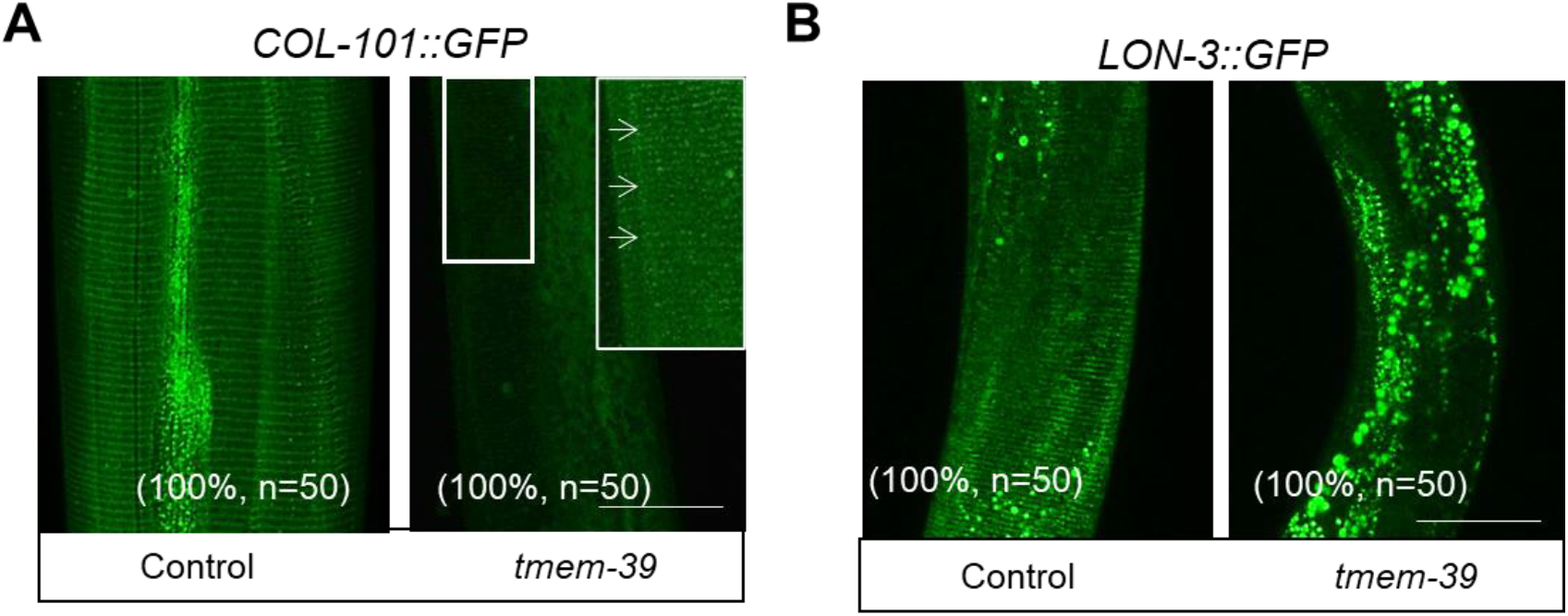
*tmem-39* RNAi knock-down in cuticle collagen translational fluorescent reporters. (A-B) Exemplar fluorescence images showing translational reporters for (A) *col-101 and* (B*) lon-3.* In wild-type animals at 20 °C (n = 3-4 for each reporters). Scale bars: 20 µm.

**S4 Fig.**
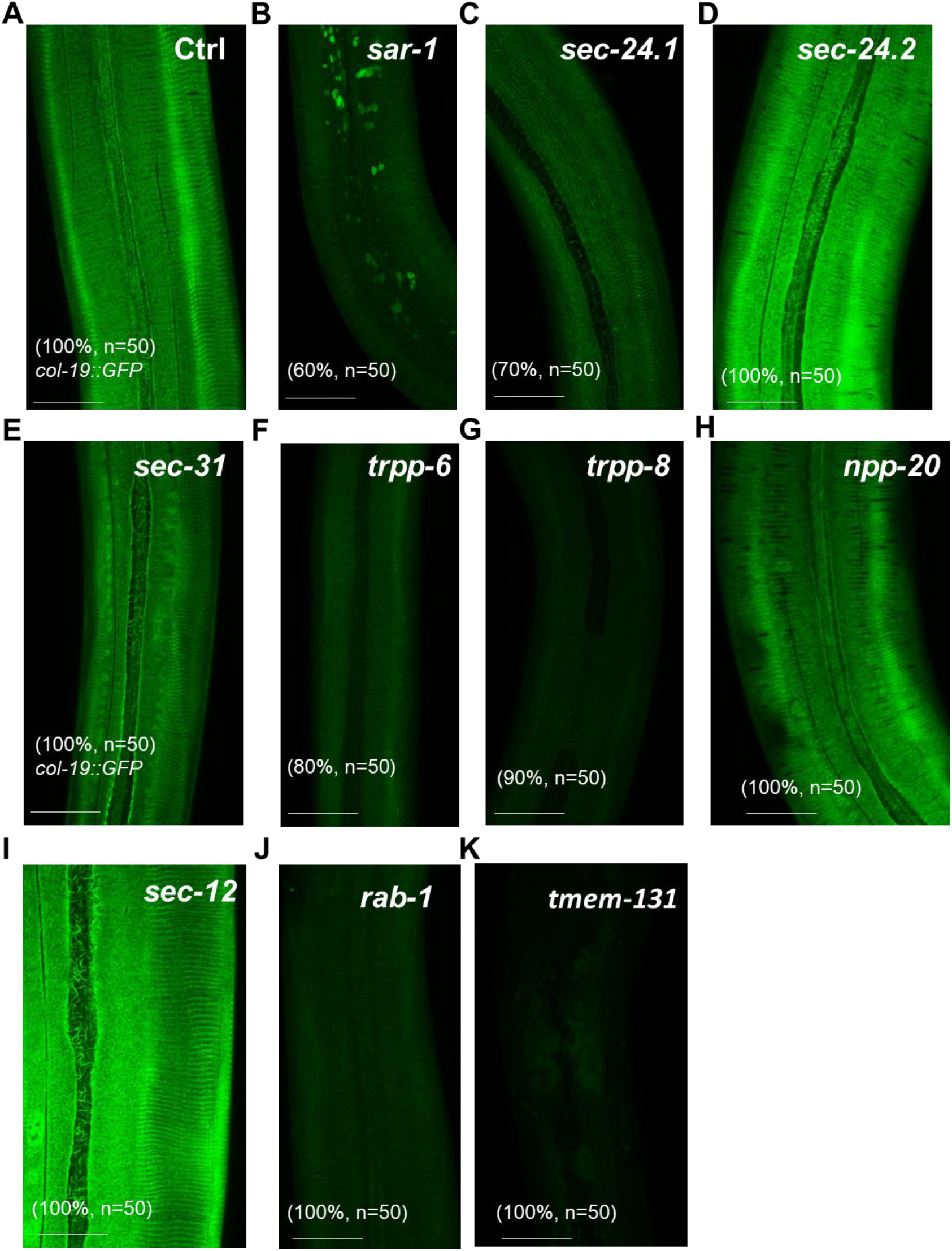
RNAi knock-down COP II component genes screen for collagen production defection. (A-J) Exemplar fluorescence images of *col-19* translational reporter for (A) control, (B) *sar-1*, (C) *sec-24.1*, (D) *sec-24.2*, (E) *sec-31*, (F) *trpp-6*, (G) *trpp-8*, (H) *npp-20*, (I) *sec-12,* (J) *rab-1* and (K) *tmem-131* RNAi in wild-type animals at 20 °C. Scale bars: 20 µm.

**S5 Fig.**
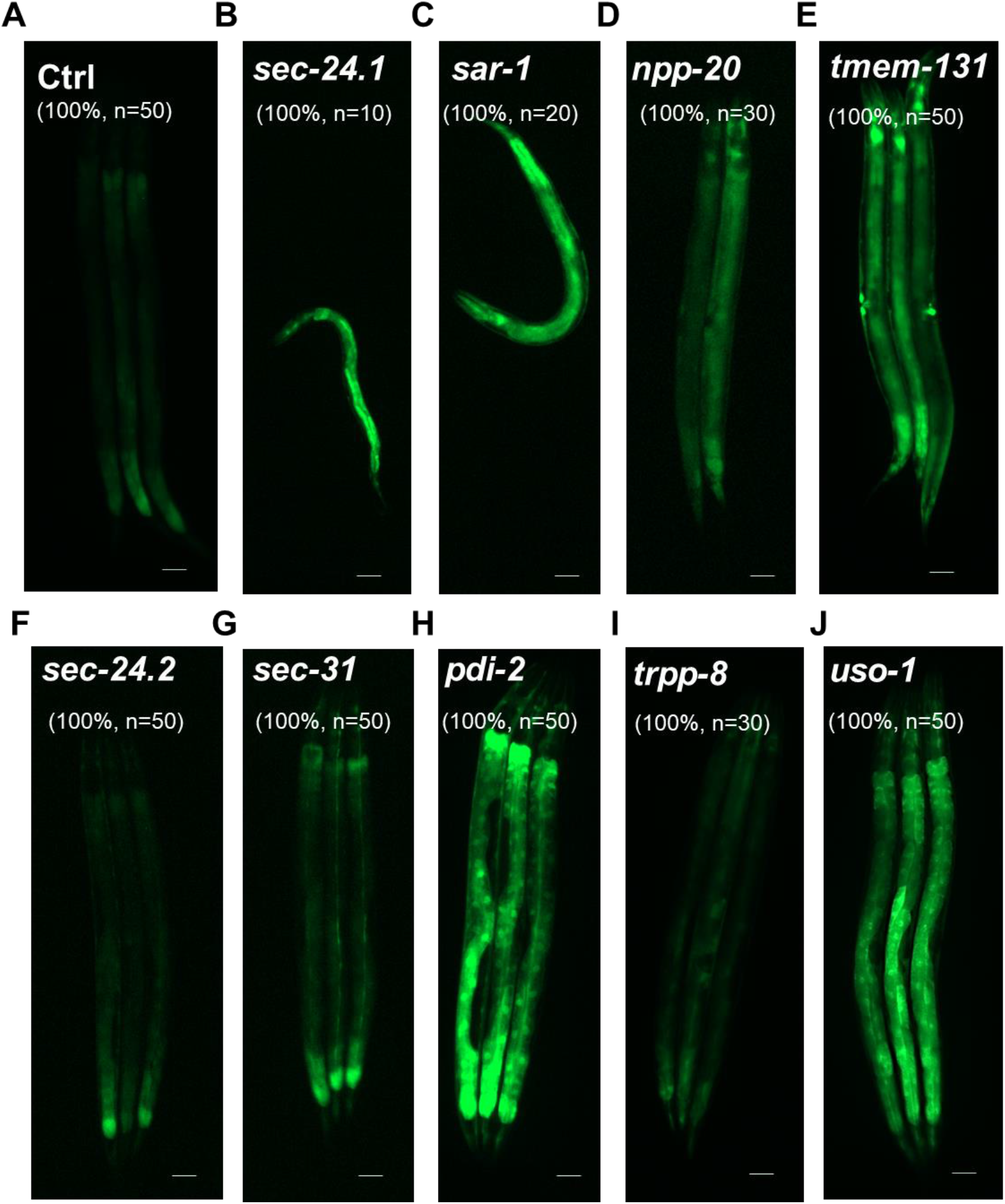
RNAi knock-down COPII component genes for screen genes involved in ER stress response. (A-J) Exemplar fluorescence images of *hsp-4*p::GFP transcriptional reporter for (A) control, (B) *sec-24.1*, (C) *sar-1*, (D) *npp-20*, (E) *tmem-131*, (F) *sec-24.2*, (G) *sec-31*, (H) *pdi-2*, (I) *trpp-8* and (J) *uso-1*RNAi in wild-type animals at 20 °C. Scale bars: 20 µm.

**S6 Fig.**
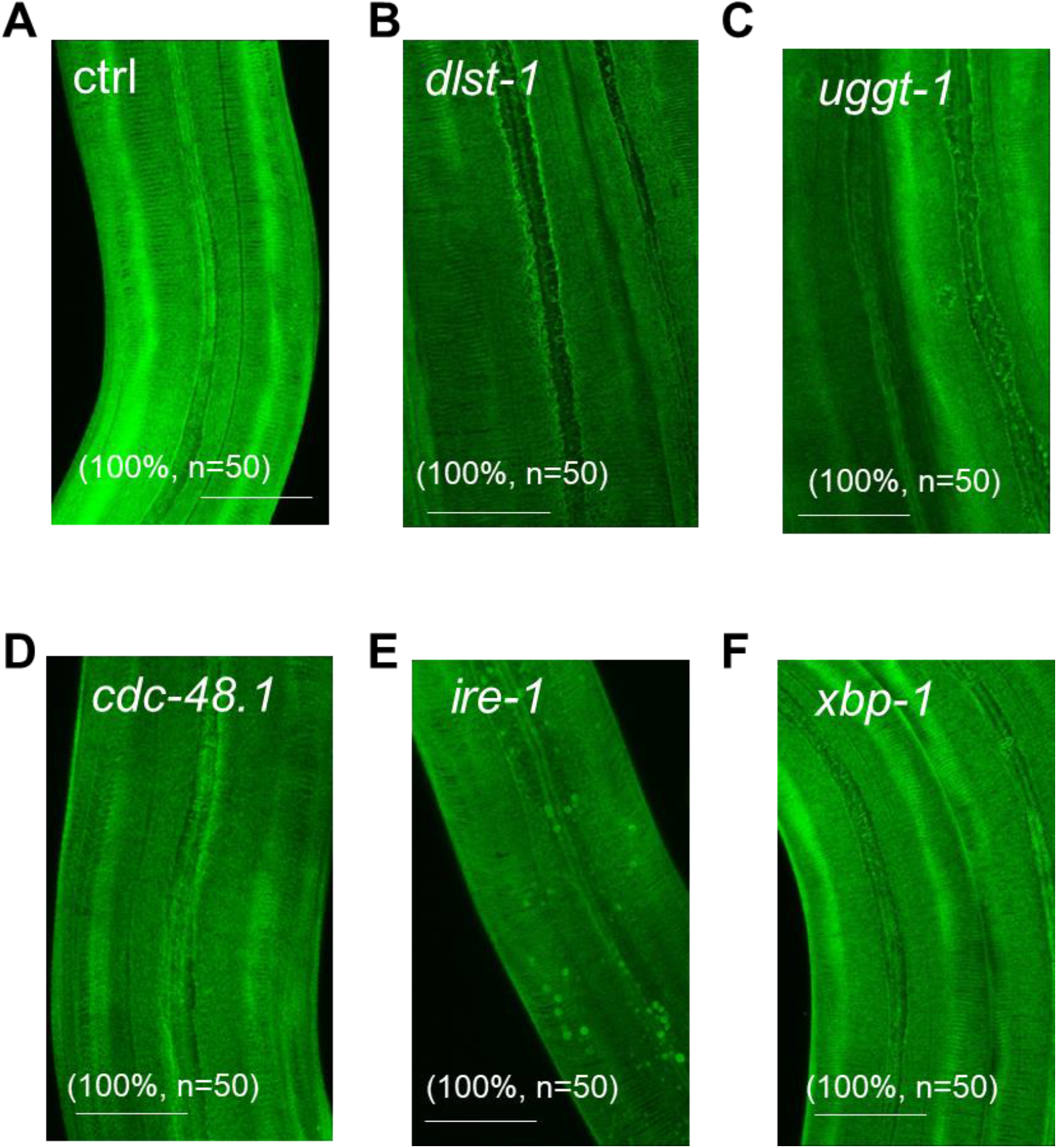
RNAi knock-down ER stress response genes in screen for collagen production defection. (A-F) Exemplar fluorescence images of *col-19* translational reporter for (A) control, (B) *dlst-1*, (C) *uggt-1*, (D) cdc-48.1, (E) *ire-1* and (F) *xbp-1* in wild-type animals at 20 °C. Scale bars: 20 µm.

**S7 Fig.**
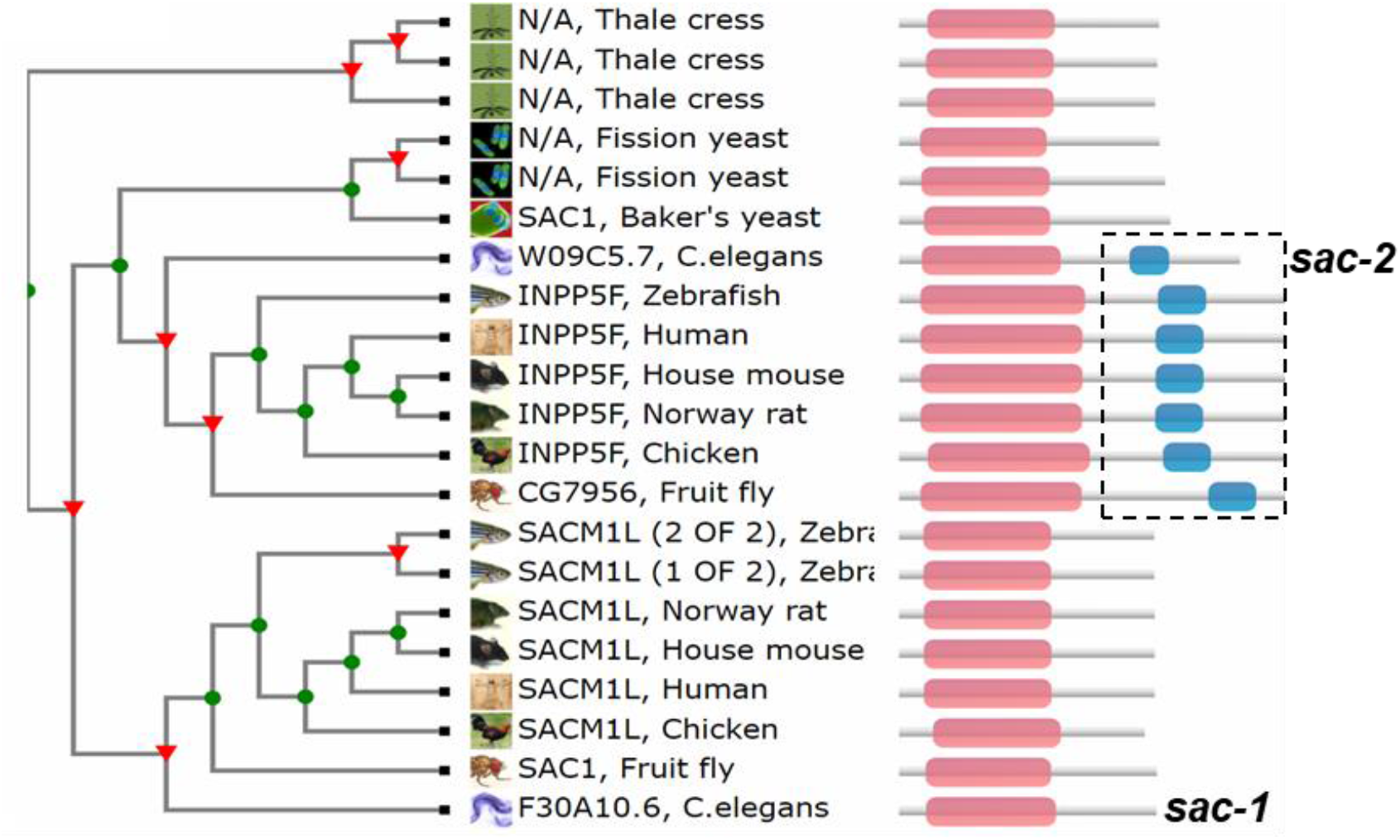
Cladogram showing conservation of the SAC1 protein sequences throughout evolution. Cladogram of phylogenetic tree for the SAC1 protein family from major representative Eukaryotic species (adapted from www.treefam.org). Domain architectures of SAC1 family proteins (right).

**S1 Table.**
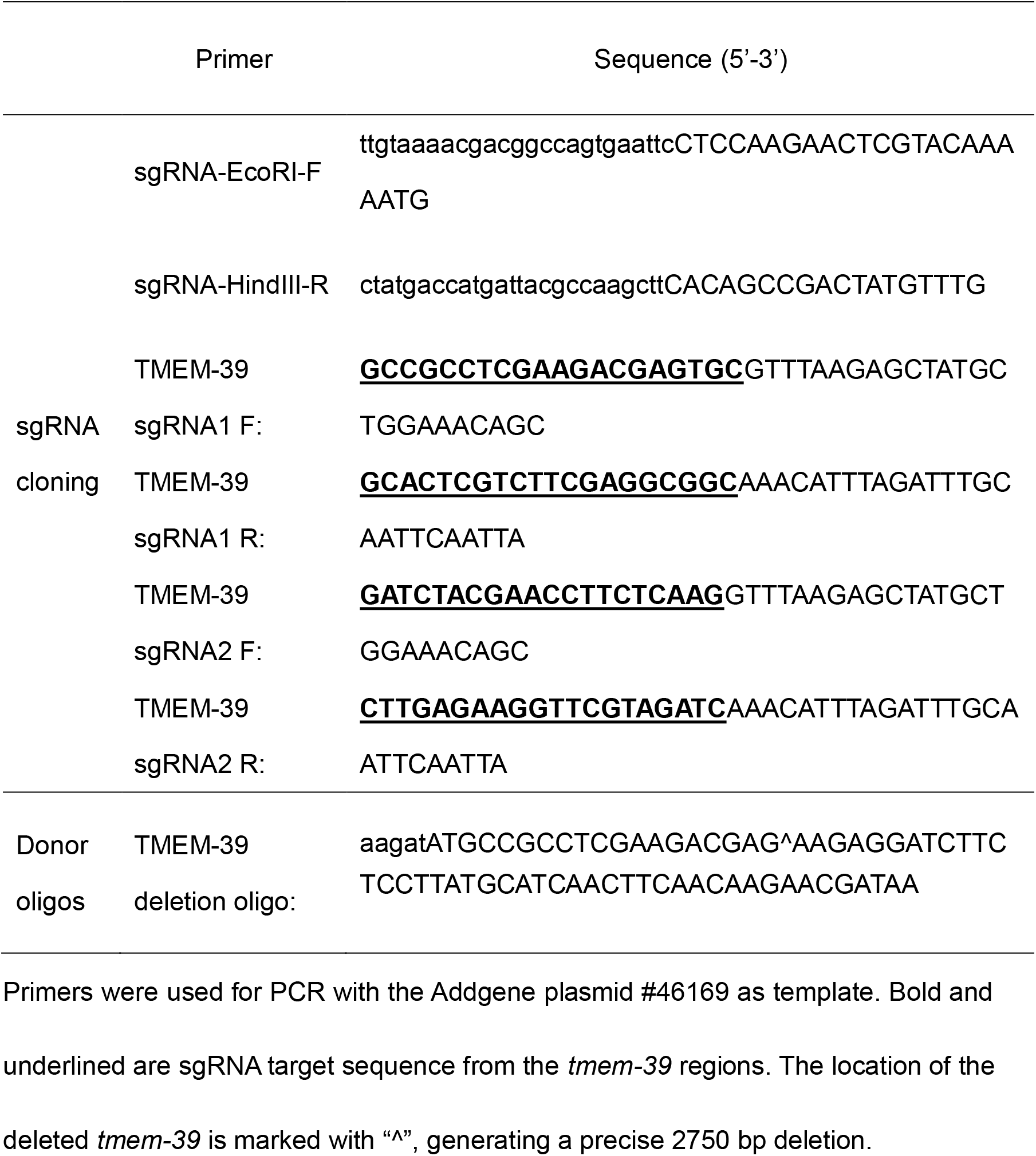
Primers and oligos used in genomic editing.

**S2 Table.**
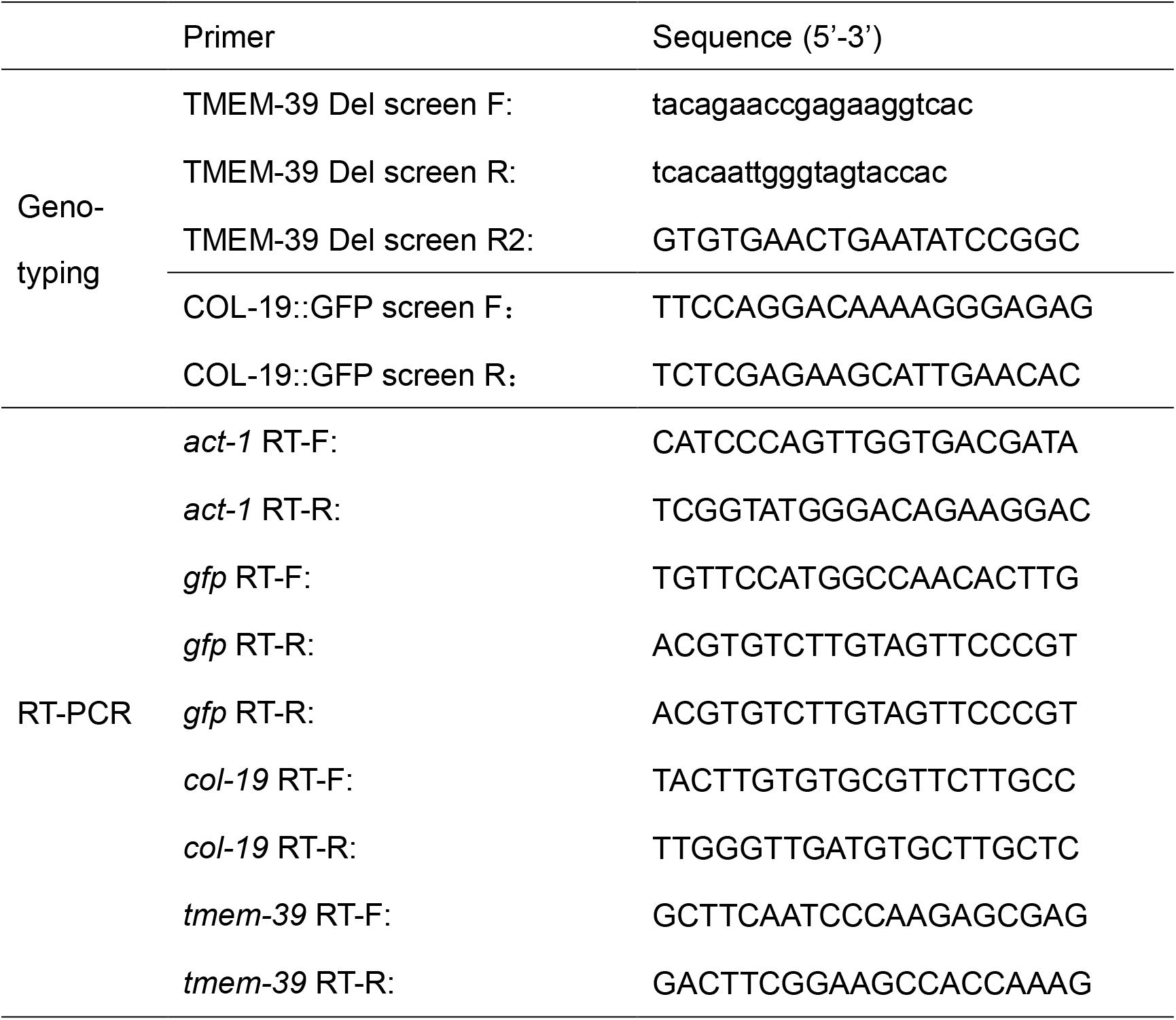
Primers in Genotyping and RT-PCR.

**S3 Table.** Reporters examined in phenotypic screen for *tmem-39* RNAi.

| Genotype | Reporter | Control | *tmem-39* |
| --- | --- | --- | --- |
| *hsp-4p::GFP* | *hsp-4* transcriptional | * | **** |
| *col-19p::col-19::GFP* | *col-19* translational | **** | * |
| *cat-1p::cat-1::GFP* | *cat-1* translational | ** | ** |
| *eff-1p::eff-1::GFP* | *eff-1* translational | * | ** |
| *emb-9p::emb-9::mCherry* | *emb-9* translational | *** | *** |
| *lrp-1p::lrp-1::GFP* | *lrp-1* translational | ** | ** |
| *fat-7p::fat-7::GFP* | *fat-7* translational | *** | *** |
| *gna-1p::GFP::gna-1* | *gna-1* translational | ** | ** |
| *him-4p::GFP::him-4* | *him-4* translational | ** | *** |
| *hmr-1p::hmr-1::GFP* | *hmr-1* translational | *** | *** |
| *lgg-1p::mCherry::GFP::lgg-1* | *lgg-1* translational | ** | ** |
| *lim-7p::ced-1::GFP* | *ced-1* translational | ** | ** |
| *myo-3p::EGFP::wrk-1* | *wrk-1* translational | *** | **** |
| *nhx-2p::cpl-1::YFP* | *cpl-1* translational | ** | *** |
| *nhx-2p::cpl-1(W32A Y35A)::YFP* | *cpl-1(W32A Y35A)* translational | *** | *** |
| *nhx-2p::ubiquitin-V::mCherry* | ubiquitin-V proteins | *** | *** |
| *rpl-28p::F23H12.5::mCherry* | F23H12.5 translational | **** | **** |
| *rpl-28p::T19D2.1::mCherry* | T19D2.1 translational | **** | **** |
| *rpl-28p::Y73E7A.8::mCherry* | Y73E7A.8 translational | **** | **** |
| *spon-1p::spon-1::vGFP* | *spon-1* translational | ** | ** |
| *unc-22p::egl-20::GFP* | *egl-20* translational | * | *** |
| *unc-54p::RFP::SP12* | ER membrane RFP marker | ** | *** |
| *unc-54p::mig-23::GFP* | *mig-23* translational | *** | *** |
| *vha-6p::mans::GFP* | intestinal GFP marker for the Golgi | *** | *** |

\* indicates fluorescent reporter levels under control and *tmem-39* RNAi conditions;
qualitative changes were followed up by quantitative fluorescence (n ≥ 20 biological
replicates) and Western blot for verification.

**S4 Table.**
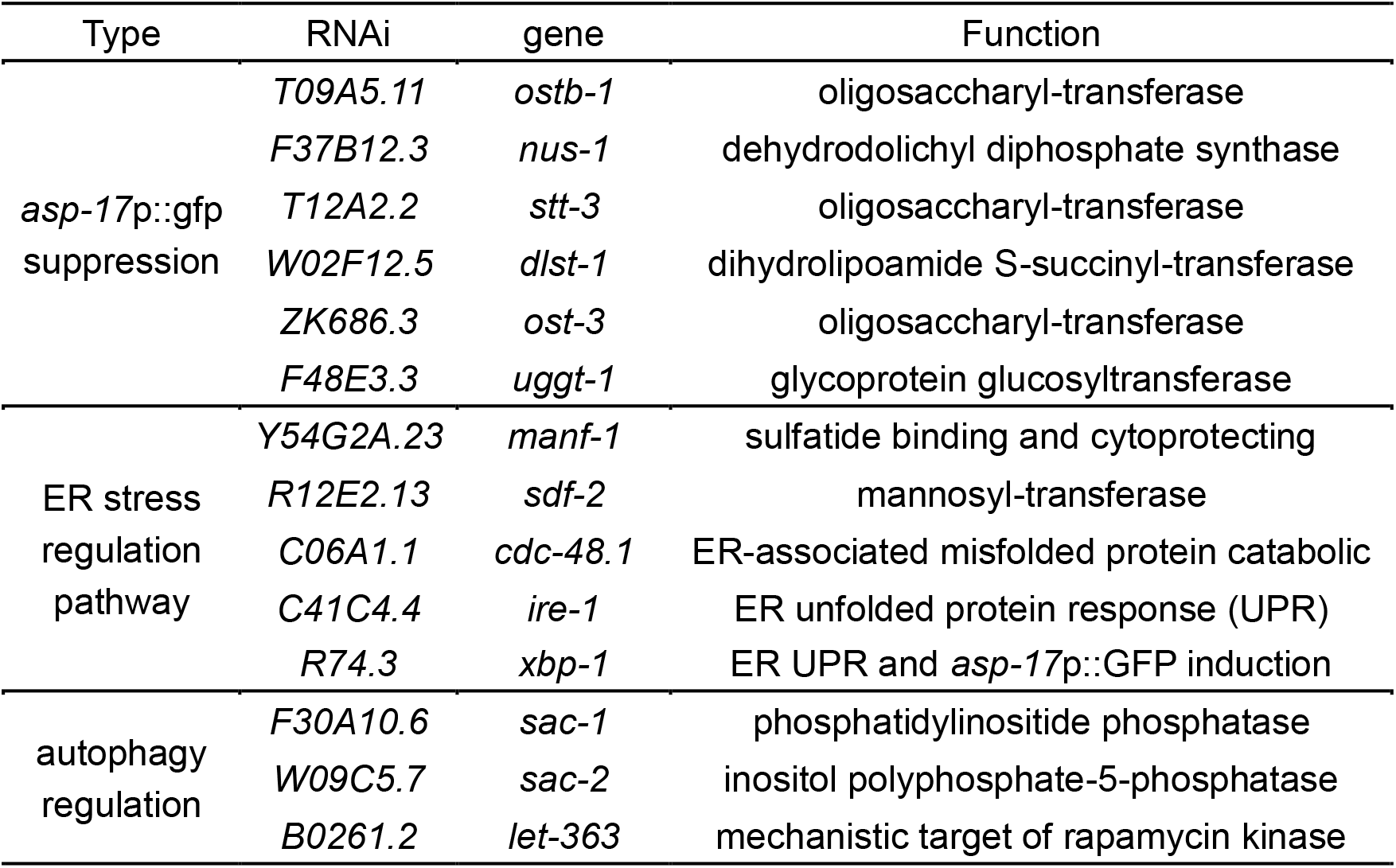
RNAi phenotypic analysis of genes for collagen secretion.

